# CREB5 promotes immunotherapy resistance via tumor-intrinsic collagen matrix deposition

**DOI:** 10.1101/2025.04.22.649109

**Authors:** Payal Tiwari, Kayla J. Colvin, Sarah Y. Kim, Ashwin V. Kammula, Jordan M Chinai, Nuno Alfaiate, Or-Yam Revach, Erin Kistler, Seoyun Heo, Seth Anderson, Sydney M. Moyer, Zhengyang Sun, Angelina M. Cicerchia, Claire A. Palin, Maulik Vyas, Cun Lan Chuong, Jessica A Talamas, Moshe Sade-Feldman, Samuel S. Freeman, Genevieve M. Boland, Kathleen B. Yates, Nir Hacohen, Shadmehr Demehri, John G. Doench, Russell W. Jenkins, William C. Hahn, Robert T. Manguso

## Abstract

Treatment with immune checkpoint inhibitors induces remarkable clinical responses in several cancer types. However, most cancer patients fail to respond to immunotherapy, and patients who initially respond often exhibit acquired resistance. Understanding the universe of immune evasion strategies will enable design of more effective immunotherapies. Here, we identify genes that drive immune evasion using genome-scale *in vivo* CRISPR gain-of-function screens in tumors treated with anti-PD-1 antibodies and found that the transcription factor CREB5 drives immune checkpoint blockade resistance. Using transcriptional profiling and functional studies, we show that CREB5 promotes a mesenchymal-like phenotype in melanoma characterized by upregulation of extracellular matrix genes including collagen and collagen-stabilizing factors. Using engineered tumor models and knockout mice, we found that immunotherapy resistance is functionally mediated by tumor-intrinsic collagen deposition. Collagen is the major ligand for the inhibitory receptor LAIR1, broadly expressed on T cells, B cells, NK cells, and myeloid cells. Deletion of LAIR1 in mice or overexpression of the decoy receptor LAIR2 in tumors abrogated the resistance induced by CREB5 overexpression, demonstrating that collagen-LAIR1 inhibitory signaling drives resistance to immune checkpoint inhibitors. These observations define a transcriptional program that remodels the tumor microenvironment to promote immunotherapy resistance via extracellular matrix deposition and indicates that targeting this pathway may enhance immunotherapy efficacy.

**One-Sentence Summary:** *In vivo* gain-of-function screening in immunotherapy-treated mice reveals a transcription factor, *Creb5*, that drives the mesenchymal state in melanoma and facilitates immune escape by promoting tumor-intrinsic collagen matrix deposition.

## Main

Cancer immunotherapy with immune checkpoint blockade (ICB) induces prolonged responses in a subset of cancer patients. However, most cancer patients do not respond to immunotherapy, and patients who initially respond often acquire resistance. Tumor-intrinsic immune evasion mechanisms are a major driver of resistance to immunotherapy and identifying and targeting these mechanisms may enable effective combination therapies. Here, we use a pooled gain-of-function screening approach with CRISPR activation to identify immune evasion programs in melanoma treated with immune checkpoint blockade.

### In vivo CRISPR activation screen identifies overexpression of CREB5 as novel mechanism of ICB resistance

To identify novel mechanisms of resistance to immune checkpoint blockade (ICB), we adapted an *in vivo* pooled genetic screening approach to perform a gain-of-function screen using the PP7-p65-HSF1 CRISPR activation system (CRISPRa), where the addition of MS2 loops into the tracrRNA sequence facilitates the recruitment of the transcription factors P65 and HSF1 to enhance gene activation (**Fig. 1A)**(*1*). To validate this strategy, we engineered the B16 mouse melanoma line to express catalytically inactivated mutant Cas9 fused to VP64 (dCas9)(*2*) (hereafter B16-dCas9) and activating single guide RNAs (asgRNAs) targeting the MHC-I molecule *H2-K1* and the “don’t eat me’’ signal *Cd47,* which inhibits phagocytosis of tumor cells by macrophages. We picked these targets because their protein expression on the surface can be evaluated using flow cytometry to assess CRISPRa activation, and both are known regulators of ICB response. We hypothesized that overexpression (OE) of *H2-K1* or *Cd47* would promote sensitivity or resistance to ICB, respectively, allowing assessment of whether CRISPRa-mediated gene induction of these targets alters the response to ICB *in vivo*. We confirmed CRISPRa-mediated OE of both genes using flow cytometry (**Fig. 1B**). Notably, we observed substantial variation in the levels of OE between different asgRNAs targeting the same gene (**Fig. S1A**). To determine whether we could alter the response to ICB using CRISPRa, we implanted B16-dCas9 cells and either control, *Cd47,* or *H2-K1* asgRNA into both untreated C57BL/6 wild-type (WT) mice and mice treated with anti-PD-1 and GVAX (GM-CSF-secreting, irradiated B16 cell vaccine). While OE of these genes did not significantly impact tumor growth in untreated mice (**Fig. S1B**), we observed that OE of *H2-K1* led to tumor regression while *Cd47* OE increased tumor growth in animals treated with immunotherapy (**Fig. 1C**). These results validated the CRISPR activation system as a method for testing effects of gene OE on immunotherapy response *in vivo*.

**Fig 1:**
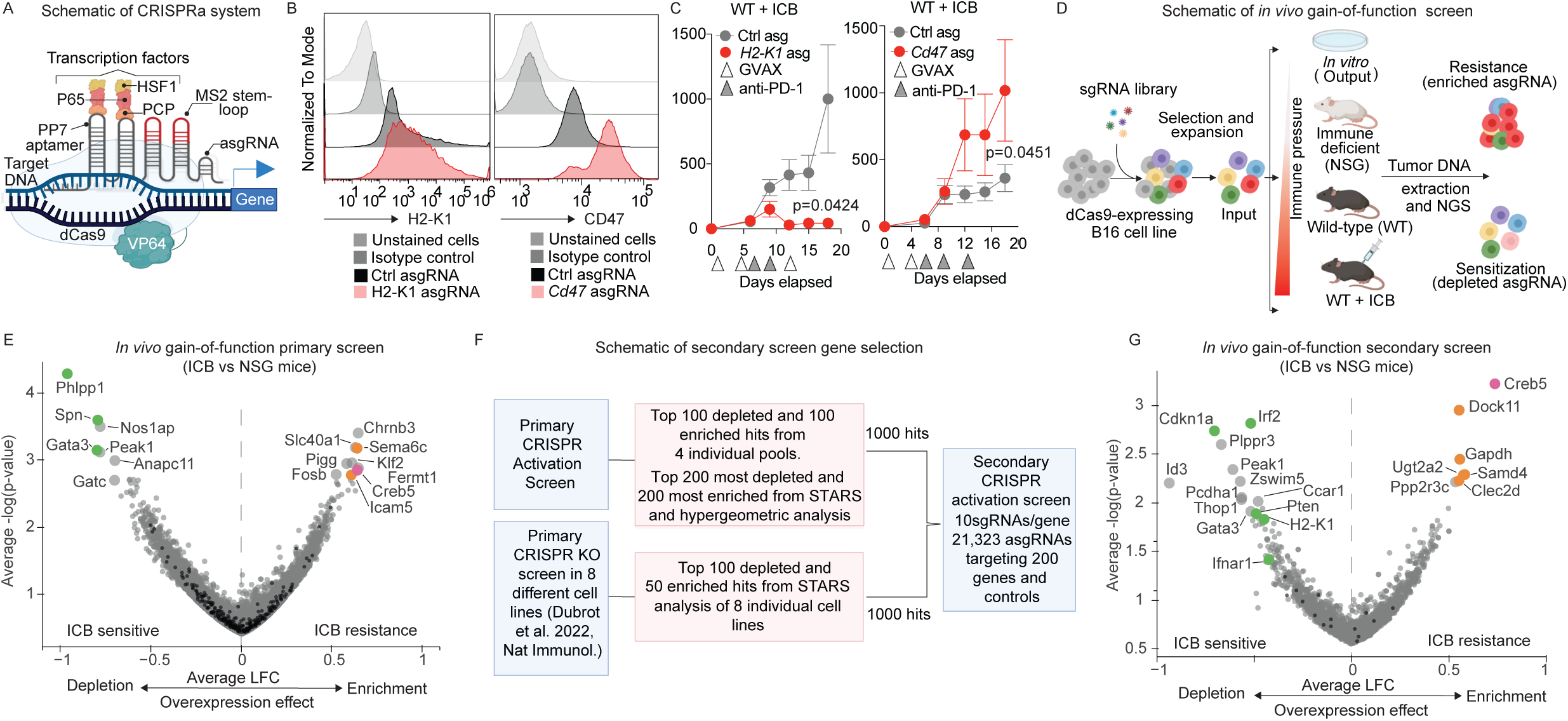
*In vivo* CRISPR activation screen identifies overexpression of Creb5 as a novel mechanism of ICB resistance. A. Schematic of CRISPR activation system. B. Histograms showing *H2-K1* and *CD47* staining in B16-dCas9 cells. C. Tumor growth over time for ICB treated mice for *H2-K1* asgRNA tumors (red) relative to control (gray) (on left) and *Cd47* asgRNA B16-dCas9 tumors (red) relative to control (gray) (on right). D. Schematic of in vivo pooled gain-of-function screen. E. Volcano plot of primary CRISPR activation screen hits. Genes marked in color are known immune regulators or novel hits of interest. Depleted targets of interest are marked in green and enriched targets of interest are marked in orange and *Creb5* is marked in Red to highlight the top enriched gene across both screens. F. Schematic for secondary screen gene selection. G. Volcano plot of secondary CRISPR activation screen hits. Genes marked in color are known immune regulators or novel hits of interest. Depleted targets of interest are marked in green and enriched targets of interest are marked in orange and *Creb5* is marked in Red to highlight the top enriched gene across both screens.

Having demonstrated the potential for CRISPR activation to identify modulators of the ICB response, we performed a genome-scale *in vivo* CRISPRa screen to identify tumor immune evasion genes (**Fig. 1D**). We designed lentiviral asgRNA libraries targeting 18,000 protein coding genes with 4 asgRNAs per gene, plus intergenic and non-targeting control guides, and divided these asgRNAs into four pools, each containing one asgRNA targeting each gene. After transduction of B16-dCas9 cells, antibiotic selection, and *in vitro* passage to allow gene activation, we transplanted the cells into three cohorts of mice: 1) WT mice treated with GVAX and an anti-PD-1 antibody to apply immune-selective pressure on the tumors (ICB); 2) untreated WT mice; and 3) immunodeficient NOD SCID IL2RG^−/−^ (NSG) mice, which lack T, B, and NK cells (**Fig. 1D**). On day 15 after implantation, we collected the tumors, isolated genomic DNA, amplified the asgRNA region, and performed sequencing to quantify asgRNA abundance. The vast majority of both target and control asgRNAs were well represented in every condition and replicates were well correlated within conditions (Pearson correlation > 0.9; **Fig. S1C-D**), confirming the quality and robustness of the screen.

To identify immune evasion genes, we compared the asgRNA representation in tumors from immunotherapy-treated WT animals to those growing in NSG mice. We identified several genes that when overexpressed either promote ICB sensitivity or resistance, including *Phlpp1*, *Spn*, and *Gata3* (depleted) and *Sema6c*, *Creb5*, and *Icam5* (enriched) (**Fig. 1E, Table S1-2**). While our primary screen identified multiple immune evasive genes, the concordance between asgRNAs targeting the same gene was weak (**Table S1-2**), consistent with the high degree of variation we observed among asgRNAs we tested individually. This is likely due to the variation in asgRNA on-target activity or the uncertainty of transcription start site (TSS) annotations for the mouse genome, potentially leading to a large number of false-negatives in our primary screening data. Therefore, to further validate candidates from the primary screen, we performed a secondary *in vivo* gain-of-function screen using a library made up of the top hits from our primary gain-of-function screen and also included the top hits from the genome-scale loss-of-function screens we previously performed to identify novel immune evasion genes(*3*). Specifically, we generated lentiviral asgRNA libraries encoding 21,323 asgRNAs targeting 2118 protein coding genes with 10 asgRNAs targeting each gene, and including 300 control guides (**Fig 1F**, see **Methods**). We performed the screen using the same conditions as our primary CRISPRa screen (Fig. 1D). Similar to the primary screen, we observed excellent guide representation and high replicate correlation, confirming the quality of the secondary screen (Pearson correlation >0.7; **Fig. S1E-F**). To confirm that hits identified from the *in vivo* conditions were not impacting growth or survival *in vitro*, and were not artifacts on engraftment into animals, we compared both the ICB-treated and NSG mouse conditions to the *in vitro* populations of cells before engraftment into animals (“input library”). As expected, immune-specific regulators such as H2-K1 and Creb5 scored strongly in the ICB-treated animals and were not identified as strongly depleted or enriched in NSG mice (**Fig. S1G**). The secondary screen revealed both depleted and enriched gene targets with established links to immunotherapy sensitivity. Among the most depleted targets were *Irf2* and *H2-K1,* overexpression of which increases MHC-I presentation(*4*), and the tumor suppressor *Pten*, loss of which has been linked to immune evasion(*5*). We also observed that overexpression of the cell cycle regulator *Cdkn1a* and the type I IFN receptor *Ifnar1* caused enhanced immunotherapy sensitivity (**Fig. 1G**). Interestingly, overexpression of the cell cycle regulator *Ccar1* also enhanced ICB sensitivity, which is concordant with our previous finding that loss of *Ccar1* caused ICB resistance in B16 melanoma in vivo CRISPR screens(*3*). Among the enriched gene targets which led to ICB resistance when overexpressed, we identified *Clec2d*, the ligand for the CD161 receptor expressed on activated CD8+ T cells and NK cells, which has been linked to immune evasion in brain tumors(*6*) **(Fig. 1G, Table S3-4**). We also identified many additional genes not previously implicated in response or resistance to immunotherapy, such as *Dock11*, *Gapdh*, *Ugt2a2*, and *Samd4,* and *Creb5* (**Fig. 1G**). For the genes included in the secondary screening library, we directly compared their fold change values in the ICB-treated vs NSG mouse comparison from the primary and secondary screens (**S1H**). While the majority of non-scoring gene targets showed no correlation between the two screens, as expected, we observed concordance among many of the top enriched and depleted gene targets between the two screens, including for *Peak1*, *Irf2*, *Gata3*, *Spn*, *Cdkn1a*, *Thop1*, *Gapdh*, *Dock11*, *Luzp1*, *Samd4*, and *Creb5* (**Fig. S1G**). We found *Creb5* among the most strongly enriched candidates from both screens and chose to focus on understanding the role of CREB5 in regulating tumor-intrinsic resistance to ICB.

### CREB5 promotes ICB resistance in melanoma cells

We generated two asgRNAs targeting *Creb5* and confirmed increased expression of both *Creb5* transcript and protein (**Fig. 2A**). Since *Creb5* is a transcriptional factor, we confirmed increased expression of CREB5 protein in the nucleus of cancer cells from tumor tissue (**Fig. 2B**). To validate that OE of *Creb5* promotes ICB resistance, we performed an *in vivo* competition assay where we mixed B16-dCas9 cells expressing one of two control (one intergenic-targeting and one non-targeting control asgRNA) or *Creb5* asgRNAs (total of 4 asgRNA populations, each comprising 25% of the total mixture) and injected them into NSG mice or WT mice with or without ICB treatment (**Fig. 2C**). On day 15, we collected genomic DNA (gDNA) from the resulting tumors and used sequencing to compare the asgRNA representation in tumors from ICB-treated mice to untreated WT or NSG mice. Both *Creb5* asgRNAs were overrepresented relative to control asgRNAs (average of 2 control asgRNAs) in ICB-treated mice compared to either untreated mice or tumors in NSG mice (**Fig. 2D, S2A-B**). Thus, *Creb5*-OE B16 cells exhibit a selective advantage in ICB-treated mice, suggesting that CREB5 promotes immunotherapy resistance. To further validate these findings, we implanted single populations of control asgRNA- or *Creb5* asgRNA-expressing B16-dCas9 cells into WT mice with or without ICB treatment. We found that *Creb5*-OE tumors were significantly more resistant to immunotherapy with a reduced survival rate relative to control tumors but had no growth advantage in the absence of immunotherapy (**Fig. 2E, S2C-D**). To validate these findings using an orthogonal approach, we overexpressed *Creb5* or truncated *Cd19* as a control by delivering an expression vector containing an open reading frame (ORF) downstream of the EF1a promoter to B16 cells (**Fig. S2E**) and implanted control or *Creb5* ORF B16 cells into WT mice with or without ICB treatment. As expected, *Creb5* ORF expressing tumors were more resistant to immunotherapy and exhibited a reduced survival rate compared to controls, but did not show a growth advantage in the absence of immunotherapy (**Fig. 2F**, **S2F-G)**. These observations demonstrated that *Creb5*-OE promotes immunotherapy resistance in the B16 melanoma mouse model.

**Fig 2:**
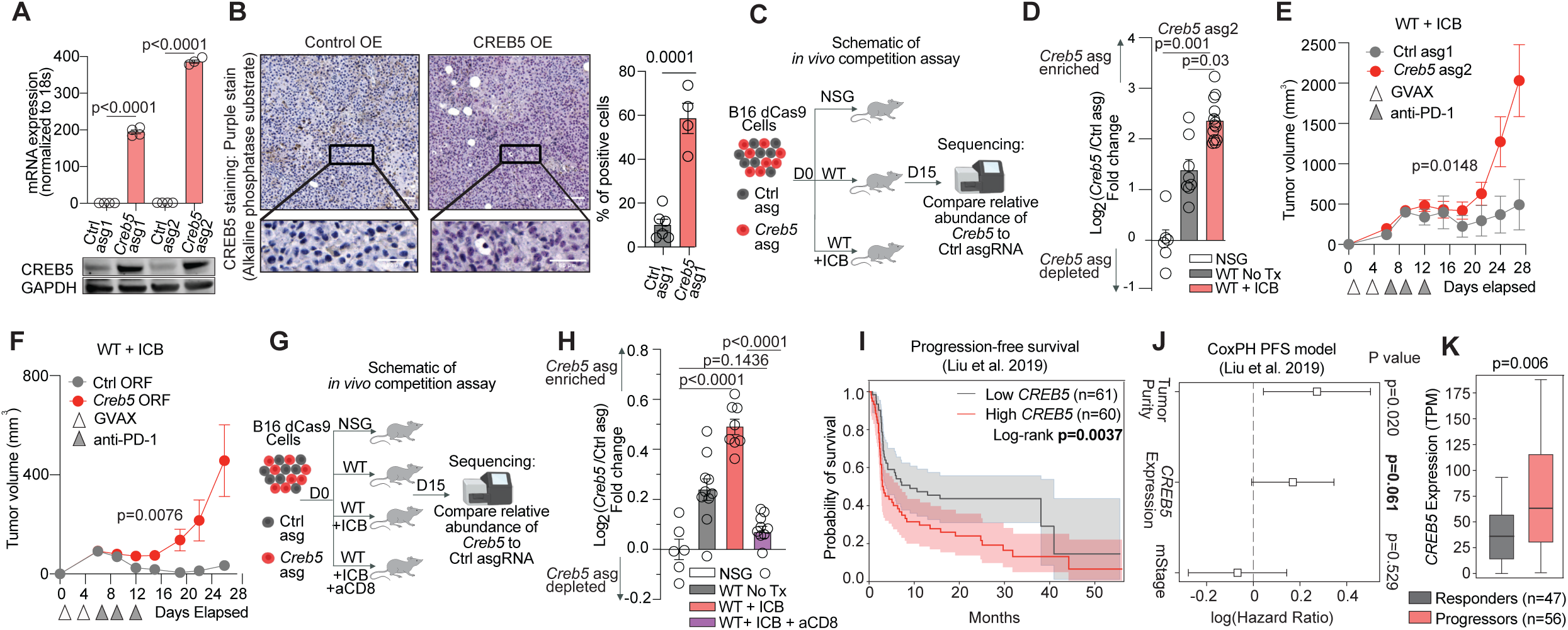
*Creb5* promotes ICB resistance in melanoma cells. A. *Creb5* transcript and protein expression in B16 control and *Creb5* asgRNA cells. B.CREB5 protein expression in nucleus (purple color) of cancer cells from tumor tissue. C. Schematic of *in vivo* competition assay. D. Results of *in vivo* competition assay showing log2 fold change for *Creb5* asgRNA relative to average of two control asgRNAs in B16 tumors. E-F. Tumor growth over time of E. *Creb5* asgRNA tumors (red) relative to control (gray) in ICB treated mice and F. *Creb5* ORF tumors (red) relative to control (gray) in ICB treated mice. G. Schematic of in vivo competition assay. H. Results of in vivo competition assay showing log2 fold change for *Creb5* asgRNA2 relative to control in B16 tumors. I. Kaplan-Meier curve showing progression-free survival for melanoma patients treated with anti-PD-1 with either high (red) or low (grey) *CREB5* expression. Low and high cohorts are separated by median *CREB5* expression. P-value from log-rank test. J. Cox proportional-hazards model evaluating the relationship between tumor purity, *CREB5* expression, and metastatic stage (mStage). All variables are z-scored prior to fitting the model. 95% confidence interval indicated. K. *CREB5* expression (TPM) in tumors from anti-PD-1-treated patients categorized as either responders (grey) or progressors (red). Difference in means calculated by Mann-Whitney U test.

Next, we tested whether ICB resistance mediated by *Creb5*-OE requires CD8^+^ T cells. We performed additional competition experiments in which we mixed B16-dCas9 cells expressing one of two control or two *Creb5* asgRNAs at equal ratios (total of 4 asgRNA populations, each comprising 25% of the total mixture) and injected them into mice in the presence or absence of ICB; a cohort of ICB-treated mice also received anti-CD8a antibody to deplete CD8^+^ T cells (**Fig. 2G**). CD8^+^ T cell depletion completely inhibited the effect of ICB on tumor growth (**Fig. S2H**), consistent with the requirement of CD8^+^ T cells for ICB efficacy. As expected, *Creb5*-OE cells showed a significant selective advantage in ICB treated tumors (**Fig. 2H, S2I**). However, the competitive advantage for *Creb5*-OE cells was completely lost in mice treated with ICB and anti-CD8a antibody (**Fig. 2H, S2I**), suggesting that direct and/or indirect selective pressure from CD8^+^ T cells favors outgrowth of *Creb5*-OE cells.

We extended these findings to a second mouse tumor model using the YUMMER (Braf^V600E^/PTEN^−/−^) mouse melanoma cell line. YUMMER cells endogenously express high levels of CREB5 relative to B16 cells (**Fig. S2J**), and we tested the effect of loss or gain of *Creb5* expression on ICB efficacy. We delivered sgRNAs targeting *Creb5* or non-targeting control in YUMMER cells engineered to express functional Cas9 nickase (YUMMER-Cas9) and observed a reduction in *Creb5* transcript and protein (**Fig. S2K**). We used the SCAR vector system(*7*) to eliminate Cas9 and selection marker proteins, as these proteins are immunogenic in YUMMER cells when injected into mice. To determine whether CREB5 mediates resistance to ICB in this model, we performed an *in vivo* competition experiment in which we mixed YUMMER-Cas9 cells expressing control sgRNA or *Creb5* sgRNA at an equal ratio, injected them into NSG mice or WT mice treated with anti-PD-1, and harvested tumors for analysis on day 15 following implantation. We compared sgRNA abundance in tumors from anti-PD-1-treated WT mice to tumors growing in NSG mice and observed that *Creb5* sgRNA was depleted relative to control sgRNA (**Fig. S2L**), indicating that loss of *Creb5* in YUMMER tumors renders them more sensitive to anti-PD-1. To evaluate whether increased expression of *Creb5* enhanced resistance to anti-PD-1, we introduced *Creb5* or control asgRNAs in CRISPRa-engineered YUMMER-dCas9 cells and observed increased CREB5 expression over baseline (**Fig. S2M**). When we mixed these cells at equal ratios and implanted them into WT mice treated with ICB or NSG mice, we found that *Creb5* asgRNA was enriched relative to control asgRNA in tumors isolated from treated mice compared to NSG mice (**Fig. S2N**), demonstrating that *Creb5*-OE YUMMER cells have a fitness advantage in anti-PD-1-treated mice. These observations demonstrate that *Creb5*-OE promotes immunotherapy resistance, while loss of *Creb5* enhances immune sensitivity in melanoma. Thus, *Creb5* expression in tumor cells is sufficient to induce resistance to immunotherapy.

Finally, to examine the link between CREB5 expression and immunotherapy resistance in human cancer, we analyzed gene expression profiling of human melanoma samples derived from a cohort of 121 patients treated with anti-PD-1(*8*). We separated patients into two groups, divided by the *CREB5* median expression in the cohort. Using a log-rank test, we observed that the *CREB5* high group exhibited significantly reduced progression-free survival (PFS) when compared to those with low *CREB5* expression (p=0.0037) (**Fig. 2I**). We then performed a Cox proportional-hazards multivariate analysis and found that the correlation between CREB5 expression and worse PFS trended toward significance (p=0.061) when accounting for differences in tumor purity and metastatic stage (**Fig. 2J**). This finding showed that association between CREB5 and worse PFS is not confounded by levels of fibroblast or immune infiltration in the tumor. In accordance with these observations, we found that patients that progressed on anti-PD-1 therapy possessed higher average *CREB5* expression relative to those that exhibited a complete or partial response to the immunotherapy (p=.00657) (**Fig. 2K**). Thus, CREB5 promotes resistance to ICB in mice, and increased expression of CREB5 is associated with poor response to ICB treatment in melanoma patients.

### CREB5 promotes expression of collagen and collagen-associated genes in cancer cells

CREB5 is a transcription factor that belongs to the CRE (cAMP response element)-binding protein family. CREB5 is essential for normal development and *Creb5*^−/−^ mice die immediately after birth(*9*). Emerging evidence suggests an important role in cancer, as CREB5 is amplified and overexpressed in several cancers, promotes proliferation and metastasis, and has been shown to enhance resistance to androgen receptor antagonists in prostate cancer(*10*, *11*). However, the relationship between these observations and ICB resistance is uncertain and the mechanism by which CREB5 promotes resistance to immunotherapy is unknown. To understand how CREB5 promotes resistance to ICB, we performed RNA-seq transcriptional profiling of control and *Creb5*-OE B16-dCas9 cells using two *Creb5* asgRNAs. We found that OE of *Creb5* caused the differential expression of ∼2,000 genes, and *Creb5*-OE cells clustered separately from control cells in both hierarchical clustering and in PCA (**Fig. 3A, Fig. S3A**). Notably, several collagens were among the most differentially upregulated genes in *Creb5*-OE cells (**Fig. 3A, Table S5**). We performed gene set enrichment analysis (GSEA) on differentially expressed transcripts using the hallmark genesets (*12*) and melanoma-specific genesets(*13*) and found significant upregulation of epithelial to mesenchymal transition (EMT) related genes and a melanoma-specific mesenchymal-like gene signature in *Creb5*-OE cells (**Fig. 3B, Table S6**). Analysis of the leading edge genes from these sets revealed that the most significantly upregulated genes driving this enrichment were collagens (*Col1a1, Col4a1, Col16a1, Col5a1, Col12a1*), collagen-stabilizing factors that promote collagen crosslinking in the ECM (*Loxl1*)(*14*), factors that prevent collagen degradation by matrix metalloproteinases (*Serpine1*), and factors that promote collagen assembly (*Fn1, Sparc, and Ecm1*)(*15–17*) (**Fig. 3C-D**). These findings were confirmed in cells expressing a second *Creb5* asgRNA, which also showed significant upregulation of mesenchymal and EMT gene sets driven by collagen and collagen-associated factors (**Fig. S3A-D, Table S7-8**). We confirmed that CREB5-OE promoted expression of collagens and collagen-stabilizing genes (C*ol1a1, Col4a1, Loxl1, Serpine1*) in both B16- and YUMMER-dCas9 cells using qPCR (**Fig. 3E, Fig S3E**; top). Furthermore, deletion of *Creb5* in Cas9-expressing versions of each model reduced the expression of collagens and collagen-stabilizing genes (**Fig. 3E, Fig S3E;** bottom).

**Fig 3:**
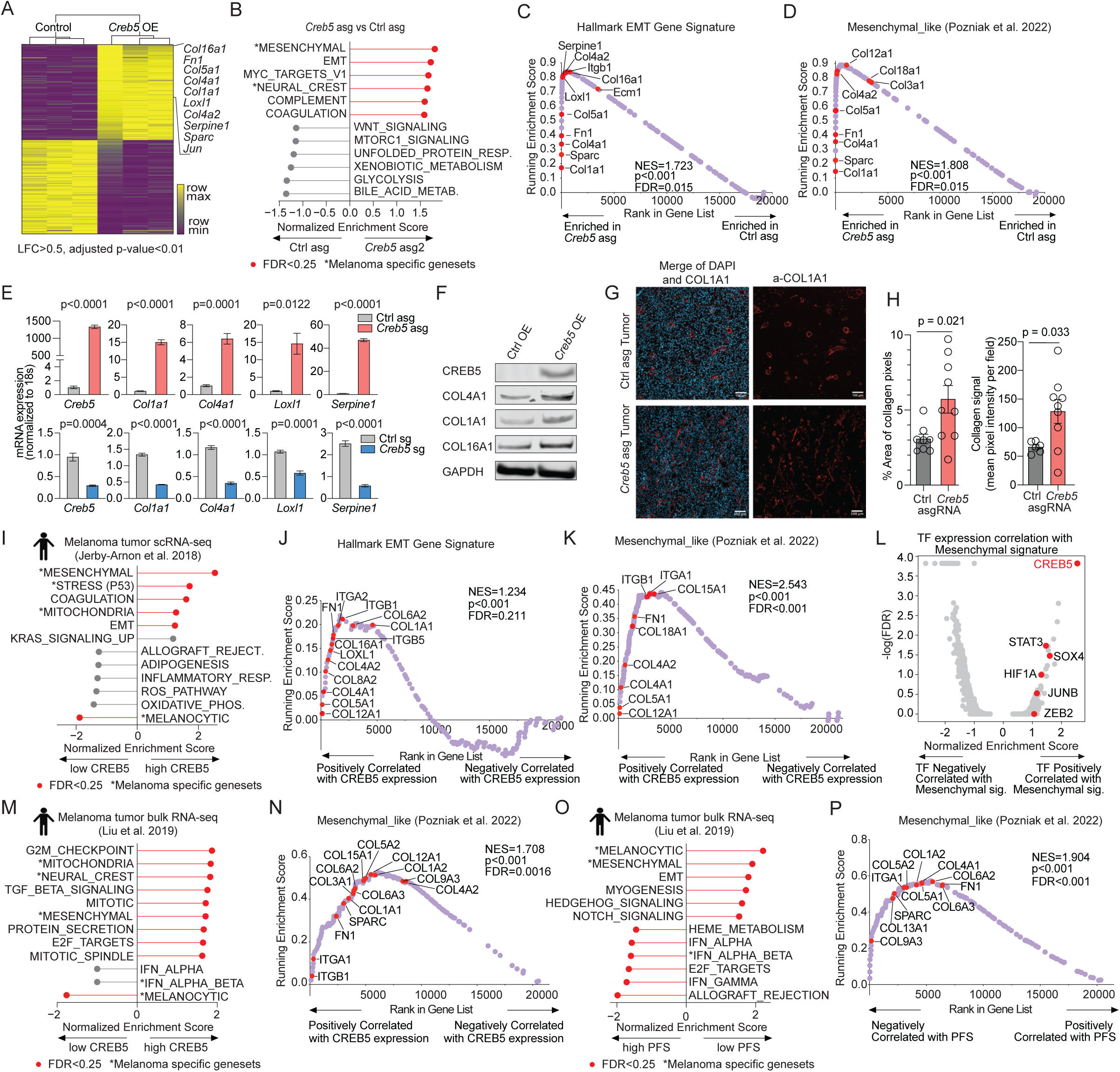
CREB5 promotes expression of collagen and collagen-associated genes in cancer cells. A. Heatmap showing hierarchical clustering and significantly differentially expressed genes between *Creb5* OE or control B16 cells. B. Bar graph showing the top 6 enriched and depleted gene sets in *Creb5* asgRNA cells relative to control B16 cells. C-D. Mountain plot showing enrichment score for the C. Hallmark EMT gene set and D. Mesenchymal-like gene set in *Creb5* asgRNA and control B16 cells. Collagen and collagen-related genes are called out. E. Transcript abundance for collagen and collagen-stabilizing factors in *Creb5* OE (top) or *Creb5* KO (bottom) B16 cells relative to control. F. Protein expression of collagen genes in B16 control and *Creb5* ORF cells. G. Immunofluorescence (IF) staining for nuclei (DAPI) and Collagen 1A1 from control (Top) or *Creb5* OE (Bottom) tumors. H. Quantification of IF staining. I. Bar graph showing the top 6 gene sets positively and negatively correlated with *CREB5* expression in malignant cells from scRNA-seq of human patient melanoma tumors. J-K. Mountain plot showing enrichment score for the J. EMT gene set and K. Mesenchymal-like gene set. Genes are ranked based on their correlation with *CREB5* expression in malignant cells from scRNA-seq of human patient melanoma tumors. Collagen and collagen-related genes are called out. L. Volcano plot showing the correlation of all transcription factors with the Mesenchymal-like gene signature. GSEA was used to quantify this relationship. Known regulators of the mesenchymal state are red. M. Bar graph showing the top 6 gene sets positively and negatively correlated with *CREB5* expression from bulk tumor RNA-seq. N. Mountain plot showing enrichment score for the Mesenchymal-like gene set. Genes are ranked based on their correlation with *CREB5* expression. O. Bar graph showing the top 6 gene sets positively and negatively correlated with progression-free survival when controlling for variation in tumor purity and metastatic stage. P. Mountain plot showing enrichment score for the Mesenchymal-like gene set. Genes are ranked based on their association with PFS calculated from Cox PH model controlling for tumor purity and metastatic stage.

To confirm that *Creb5*-OE increased collagen expression at the protein level, we performed immunoblotting of total cell lysates of either control or *Creb5*-OE B16 cells. We observed an increase in COL1A1, COL4A1, and COL16A1 in the *Creb5*-OE B16 cells relative to control cells (**Fig. 3F**). We hypothesized that CREB5 directly regulates transcription of collagen genes. Previous work in prostate cancer revealed that CREB5 binds to FOXA1 motifs (TATTTAT, TGTTTAT, TATTTAT), and we identified several FOXA1 motifs within 5kb of the promoters of *Col1a1*, *Col4a1,* and *Col16a1*(*18*). Using CUT&RUN PCR for V5-tagged CREB5 expressed in B16 cells, we observed a significant increase in the binding of CREB5 to FOXA1 motifs at the promoter of each collagen gene compared to chromosome14 region deprived of FOXA1 motifs (**Fig. S3F**). Finally, to determine whether *Creb5*-OE increased collagen deposition in tumors, we performed immunofluorescence staining for type I collagen in control or *Creb5*-OE B16 tumors and observed a marked increase in collagen I when *Creb5* was overexpressed (**Fig. 3G-H, Fig S3G**). Thus, CREB5 positively regulates collagen expression and deposition in melanoma cell lines and tumors.

Tumor-specific ECM, which exhibits higher stiffness and has a high collagen content, has been linked to a poor prognosis in several cancers by promoting immune exclusion(*19–21*). To address whether Creb5 OE alters collagen structure in B16 tumors, we performed trichrome staining in B16-dCas9 tumors expressing control, *Creb5* or *Col1a1* asgRNA. To our surprise, we failed to observe collagen fiber staining (trichrome stains collagen fibers blue) around the cancer cells in B16 tumors (**Fig. S3H**). This was unlikely due to a technical issue related to the trichrome staining, as we observed strong signals in collagen fibers at the tumor margins, which are enriched with collagen-producing stromal cells. Despite not observing positive trichrome staining which only stains collagen fibrils, when we performed immunostaining for individual collagens in these tumors we observed significantly increased collagen around cancer cells in both *Creb5* OE and *Col1a1* OE tumors relative to controls (**Fig S3I**). These observations suggest that CREB5 modulates collagen quantity rather than structure.

We next assessed whether *CREB5* promotes the expression of collagen and collagen-associated genes in human melanoma. First, we generated *CREB5*-OE and *CREB5* KO A375 melanoma cells and observed elevated expression of collagen genes in *CREB5*-OE cells and reduced expression of collagen genes in *CREB5* KO cells (**Fig. S3J**). Thus, modulation of *CREB5* expression in human melanoma cells has the expected effect on the expression of collagen factors. To determine whether *CREB5* expression is associated with collagen and collagen-associated genes in patient tumors, we examined single-cell gene expression profiling of human melanoma samples from two independent studies(*13*, *22*). In melanoma cells, we found a strong correlation between *CREB5* expression and expression of genes in the Hallmark epithelial to mesenchymal transition (EMT) pathway and the melanoma-specific mesenchymal-like gene signature using GSEA (**Fig. 3I, S3K, Table S9-10**). As we found in our murine models, analysis of the leading-edge genes from these gene sets revealed that the most significantly correlated genes driving this enrichment were collagens and collagen-associated genes (**Fig 3J-K, S3L-M**). We also performed the reciprocal analysis, where we identified the transcription factors most strongly associated with the mesenchymal-like signature. Strikingly, *CREB5* was the most significantly correlated transcription factor among all the 986 TFs detected in the dataset(*23*) (**Fig 3L, Table S11**). Similarly, in the same bulk melanoma tumor RNA-seq cohort where we showed an association between *CREB5* expression and ICB response, we also observed that *CREB5* expression significantly correlated with the mesenchymal-like signature which was again associated with upregulation of collagen genes (**Fig. 3M-N, Table S12**). We next assessed whether these signatures are associated with immunotherapy treatment outcome, as we did for *Creb5* itself. We observed that mesenchymal-like signatures, as well as the Hallmark EMT geneset, were among the most strongly associated with worse PFS with treatment with ICB, while the IFN gamma gene signature was associated with better PFS (**Fig. 3O, Table S13**). Finally, when we examined the genes driving the association between the mesenchymal-like signature and PFS, we observed many of the same collagen and collagen-associated factors (**Fig. 3P**). Thus, *CREB5* promotes a mesenchymal-like phenotype in human melanoma cells that is characterized by high levels of expression of collagen and collagen-associated genes, and expression of this mesenchymal signature is associated with poor response to ICB.

### Tumor-intrinsic collagen matrix deposition promotes ICB resistance

Given that collagen factors were tightly associated with *CREB5* expression in both mouse and human melanoma cells, we hypothesized that tumor-intrinsic collagen matrix deposition might be functionally important for resistance to ICB. Thus, we examined whether enhancing collagen expression in tumor cells is sufficient to promote resistance to immunotherapy. Specifically, we engineered B16-dCas9 cells to overexpress individual collagen genes that we found upregulated in *Creb5* OE cells: *Col1a1*, *Col4a1*, and *Col16a1* (**Fig. S4A**). We confirmed that increasing the expression of collagen genes in tumor cells increases collagen protein deposition in melanoma tumors in vivo (**Fig. S3I and S4B**). We then implanted these cells into WT mice with or without ICB treatment. Surprisingly, we found that overexpression of *Col1a1*, *Col4a1*, or *Col16a1* in tumors rendered them significantly more resistant to immunotherapy relative to controls (**Fig. 4A-C**). In the absence of immunotherapy, *Col1a1* and *Col16a1* OE tumors did not exhibit a growth advantage, while *Col4a1* OE tumors grew more aggressively compared to controls (**Fig. S4C**). These observations confirmed that increased collagen expression by cancer cells promotes ICB resistance.

**Fig 4:**
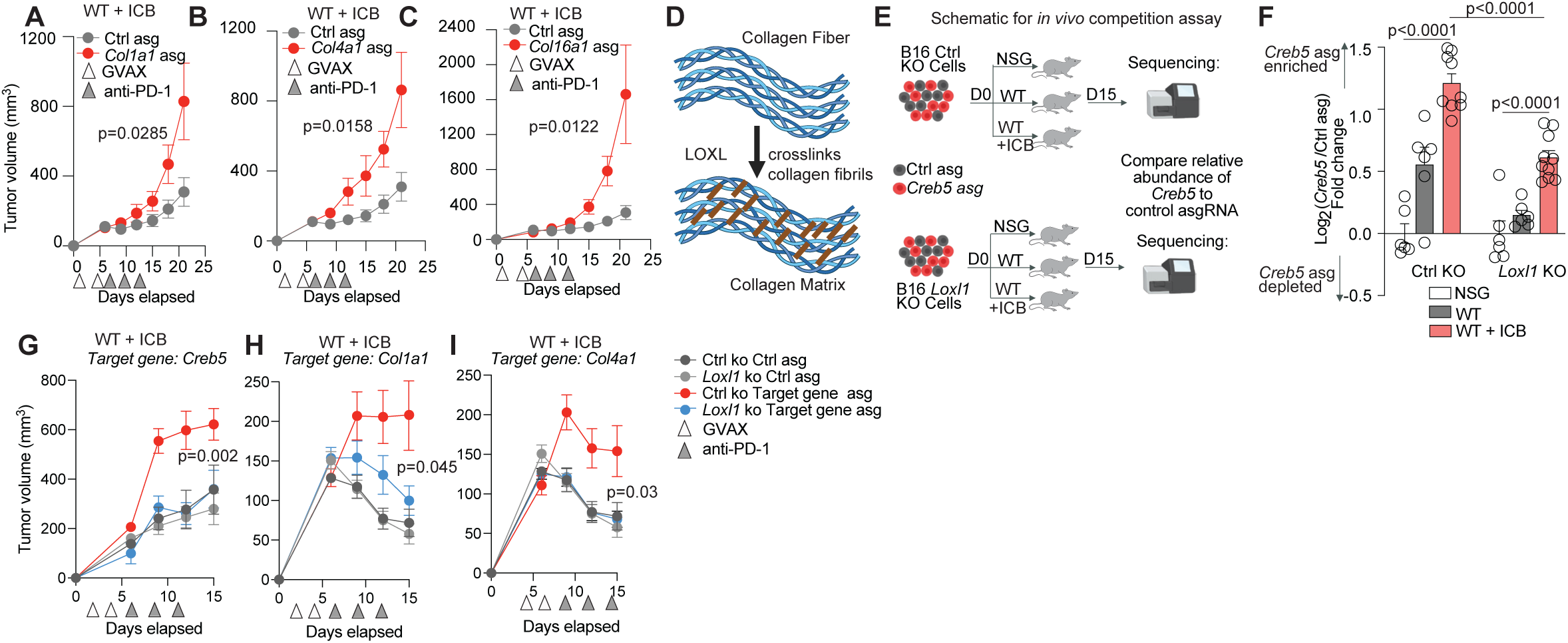
Tumor-intrinsic collagen matrix deposition promotes ICB resistance. A-D. Tumor growth over time in ICB treated mice for A. *Col1a1* OE tumors (Red) relative to control (Gray), B. *Col4a1* OE tumors (Red) relative to control (Gray), and C. *Col16a1* OE tumors (Red) relative to control (Gray). D. Schematic of LOXL-mediated collagen crosslinking. E. Schematic of *in vivo* competition assay. F. Results of *in vivo* competition assay showing log2 fold change for *Creb5* asgRNA relative to control in B16 control KO tumors (Left) or B16 *Loxl1* KO tumors (Right). G-I. Tumor growth over time in ICB treated mice for G. *Creb5* OE (Red) in *Loxl1*-deficient (Blue) or *Loxl1*-sufficient (Red) tumors relative to control OE in *Loxl1*-deficient (Gray) or *Loxl1*-sufficient (Black) tumors. H. *Col1a1* OE tumors with *Loxl1*-deficient (Blue) or *Loxl1*-sufficient (Red) tumors relative to control OE in *Loxl1*-deficient (Gray) or *Loxl1*-sufficient (Black) tumors. I. *Col4a1* OE tumors with *Loxl1*-deficient (Blue) or *Loxl1*-sufficient (Red) tumors relative to control OE in *Loxl1*-deficient (Gray) or *Loxl1*-sufficient (Black) tumors.

We next set out to determine whether collagen deposition is required for resistance phenotype caused by the OE of *Creb5*. Rather than delete individual collagen factors which could be functionally redundant, we tested this experimentally by reducing the deposition of several types of collagen through deletion of the collagen crosslinking enzyme LOXL1. LOXL1 is a member of the lysyl oxidase family of proteins that catalyzes the first step in the formation of crosslinks in collagen and promotes collagen deposition (**Fig. 4D)**(*14*), and is upregulated in *Creb5* OE cells (Fig. 3C). To investigate whether *Loxl1* deletion diminished the deposition of type I collagen in B16 melanoma tumors, we generated *Loxl1* KO or control B16-Cas9 cells (**Fig. S4D**), implanted them into WT mice, and performed immunofluorescence (IF) staining for CD31, an endothelial marker, and COL1A1. Perivascular stromal cells are a major source of collagen, and we used CD31 expression in blood vessels to distinguish which collagen was likely not produced by cancer cells. We calculated the total COL1A1 pixel area coverage and subtracted signal attributed to CD31-labeled blood vessels, and found a significant reduction in collagen deposition in *Loxl1*-deficient B16 tumors compared to controls(**Fig. S4E**). These data confirmed the key role for LOXL1 in collagen crosslinking and deposition(*24–27*).

To determine whether collagen deposition by cancer cells is required for CREB5-mediated ICB resistance, we overexpressed *Creb5* in control or *Loxl1* KO B16 cells using CRISPRa. We performed a competition assay by implanting a 1:1 mix of *Creb5* asgRNA and control asgRNA cells expressing in control or *Loxl1* KO B16-dCas9 cells into NSG mice and WT mice treated with or without GVAX and anti-PD-1 (**Fig 4E**). As expected, we found that *Creb5* asgRNA was overrepresented relative to control asgRNA in ICB-treated *Loxl1*-sufficient tumors compared to NSG mice. However, the competitive advantage for *Creb5*-OE cells was reduced by 50% relative to control in *Loxl1*-deficient tumors, suggesting that tumor-intrinsic collagen crosslinking plays a role in CREB5-mediated resistance (**Fig. 4F**). To validate these findings using in vivo tumor growth assays, we implanted B16-dCas9 cells expressing *Creb5*, *Co11a1, Col4a1*, or control asgRNA in either a control or *Loxl1* KO background into WT mice with ICB treatment. As expected, we found that *Loxl1*-sufficient tumors overexpressing *Creb5*, *Col1a1* or *Col4a1* were significantly more resistant to immunotherapy relative to controls. However, deletion of *Loxl1* completely blocked the ICB resistance phenotype for each gene (**Fig. 4G-I**). To further extend these findings we deleted *Loxl2*, another enzyme involved in crosslinking collagen in the extracellular matrix and compared this phenotype to the deletion of *Loxl1* (**Fig. S4F)**. We performed an in vivo competition assay by implanting a 1:1 mix of *Creb5* asgRNA and control asgRNA cells expressed in control or *Loxl1* KO or *Loxl2* KO B16-dCas9 cells into NSG mice and WT mice treated with GVAX and anti-PD-1 (**Fig. S4G).** As expected, we found that *Creb5* asgRNA was overrepresented relative to control asgRNA in ICB-treated control KO tumors compared to tumors from NSG mice. However, the competitive advantage for *Creb5*-OE cells was reduced by ∼50% relative to control in both *Loxl1* or *Loxl2* KO tumors, suggesting that tumor-intrinsic collagen crosslinking by both enzymes plays a role in CREB5-mediated resistance (**Fig. S4H)**. LOXL1 and LOXL2 have cellular functions beyond cross-linking collagen, and thus we cannot rule out that another function for these enzymes is related to the suppression of the CREB5-mediated resistance phenotype. However, their well-established role in collagen matrix assembly strongly suggests that this process is essential for CREB5- and collagen-mediated ICB resistance. Thus, overexpression of CREB5 and collagen genes causes resistance to ICB in melanoma and requires the intact function of lysyl oxidase enzymes.

### Creb5 overexpression promotes an immunosuppressive microenvironment

Overexpression of *Creb5* and the resulting collagen overproduction enhances cancer cell intrinsic immune evasion; however, enhanced tumor cell-derived collagen deposition could also change the composition of the tumor microenvironment and impact the frequency and spatial distribution of various immune subsets. We performed imaging by immunofluorescence microscopy to assess the impact on the tumor microenvironment. We implanted the B16 tumors with control or *Creb5* OE into anti-PD1 antibody treated mice and, as expected, observed a significant reduction in the ICB response in the *Creb5* OE tumors (**FigS5A**). We stained tumor sections for COL1A1, CD45, F4/80, CD8, CD4 and FOXP3 (**Fig. 5A-E**). To understand the distribution and infiltration of immune cells, we quantified the density of specific subsets either at the edges or at the center of tumors. In both groups, the tumor edge was characterized by a thick collagen matrix, consistent with previous studies showing that tumor-associated fibroblasts create a dense ECM at the tumor boundary(*28–30*). We observed higher collagen staining in control tumors relative to CREB5-OE tumors upon ICB treatment, which we hypothesize to be an adaptive response to denser immune infiltration and a stronger response to therapy in the control condition(*30*) (**Fig. S5A**).We did not observe a difference in CD45+ immune cells, F4/80+ macrophages, CD8+ T cells, CD4+ T cells, Foxp3-T helper (Th) cells, or Foxp3+ T regulatory (Treg) cells at the edges of CREB5-OE tumors relative to control-OE tumors (**Fig. 5B-E**), consistent with the idea that most of the collagen in this region is produced by non-tumor cells and thus not impacted by the overexpression of *Creb5*. However, in the tumor center we observed a trend towards fewer total CD45+ cells (p= 0.07), significantly fewer macrophages (p= 0.0096), a trend towards fewer CD8+ and total CD4+ cells (p=0.11; p=0.06), and significantly fewer Foxp3-CD4+ cells (p=0.04) in *Creb5* OE tumors relative to control tumors (**Fig. 5B-E**). The observation of immune composition differences in the tumor center but not the edge is consistent with the notion that tumor-intrinsic overexpression of collagen would have the greatest impact in the the region where the collagen matrix is more influenced by tumor cells rather than other cell types such as tumor-associated fibroblasts. This finding also suggests that *Creb5* OE is not creating an immune desert or exclusion phenotype but instead supporting an immune suppressive phenotype, likely through the direct engagement of immune cells with collagen.

**Fig 5:**
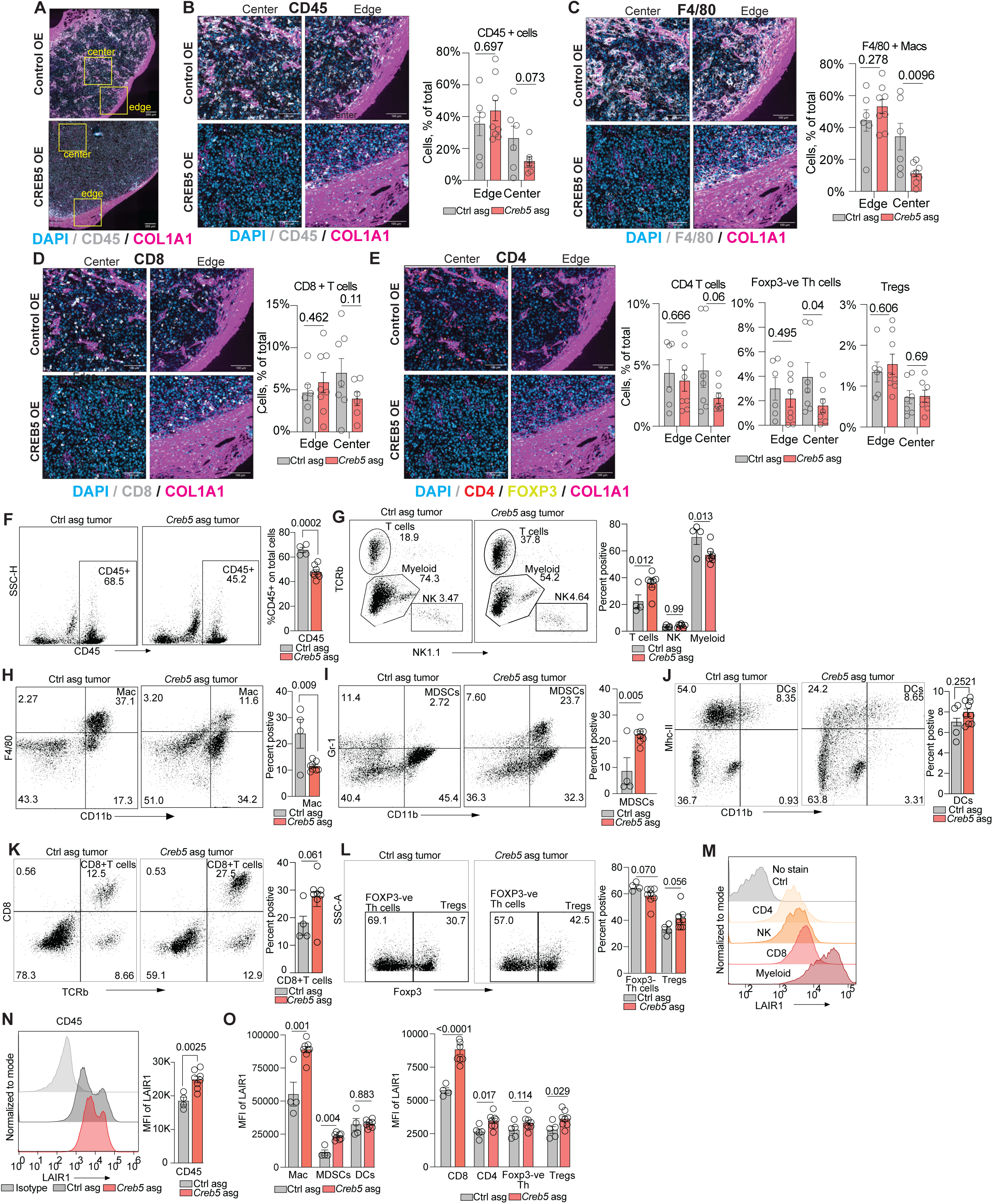
CREB5 promotes immunosuppressive tumor microenvironment. A-E. Representative images of Multiplex Immunofluorescence (IF) of paraffin embedded formalin fixed B16 tumors sections post ICB treatment with CD45, COL1A1, CD8, F4/80, CD4, FOXP3 and DAPI. All of the IF images depict the same section in which IF was performed simultaneously but analyzed in different channels. A. Zoom out picture of Control-OE tumor on top and *Creb5*-OE tumor on bottom, showing CD45 (White), Dapi (Blue), Col1a1 (Pink), with marked tumor edge and tumor center for subsequent zoom in images. B-E. Zoom in images of Control-OE tumors (Top) and *Creb5*-OE tumor (Bottom) with section of tumor center (Left) and tumor edge (Right) and quantification of total pixel per total number of cells. B. CD45 (White), Dapi (Blue), and COL1A1 (Pink). C. F4/80 (White), Dapi (Blue), and COL1A1 (Pink). D. CD8 (White), Dapi (Blue), and COL1A1 (Pink). E. CD4 (Red), FOXP3 (Yellow), Dapi (Blue), and COL1A1 (Pink). F-O. Flow cytometry analysis of TILs isolated from B16 tumors post ICB treatment. Dot plot showing control OE tumors (Left), *Creb5* OE tumors (Center), and quantification of percentage positive cells (Right). F. Dot plot showing CD45 staining on tumor cells post digestion. G. Dot plot showing TCRb and NK1.1 stains on cells post lymphocyte enrichment and gated on CD45 positive population on left and quantification of TCRb positive T cells, NK1.1 positive NK cells and TCRb and NK1.1 negative myeloid cells as percentage of total CD45 positive cells in control OE (Gray) and *Creb5* OE (Red) tumors on right. H. Dot plot showing CD11b and F4/80 stains in CD45 positive population on left and quantification of CD11b and F4/80 double positive macrophages as percentage of total CD45 positive cells in control OE (Gray) and *Creb5* OE (Red) tumors on right. I. Dot plot showing CD11b and Gr-1 stains in CD45 positive on left and quantification of CD11b and Gr-1 double positive MDSCs as percentage of total CD45 positive cells in control OE (Gray) and *Creb5* OE (Red) tumors on right. J. Dot plot showing CD11b and MHC-II stains in CD45 positive population on left and quantification of CD11b and MHC-II double positive DCs as percentage of total CD45 positive cells in control OE (Gray) and *Creb5* OE (Red) tumors on right. K. Dot plot showing TCRb and CD8 stains in CD45 positive population on left and quantification of TCRb and CD8 double positive T cells as percentage of total CD45 positive cells in control OE (Gray) and *Creb5* OE (Red) tumors on right. L. Dot plot showing Foxp3 stain in CD45, TCRb and CD4 positive population on left and quantification of Foxp3 negative T-helper cells and Foxp3 positive T regulatory cells as percentage of total CD4 T cells in control OE (Gray) and *Creb5* OE (Red) tumors on right. M. Histograms showing LAIR1 staining on CD4 positive cells (yellow), NK (Brown), CD8 (Red) and CD11b positive myeloid cells (Deep red). N. Histograms showing LAIR1 staining on CD45 positive cells and quantification of LAIR1 MFI on CD45 positive cells in control OE (Gray) and *Creb5* OE (Red) tumors on the right. O. Quantification of LAIR1 MFI on immune cells in control OE (Gray) and *Creb5* OE (Red) tumors on the right.

To further validate these findings and characterize the immune cell populations, we performed flow cytometry analysis of TILs from control and *Creb5*-OE B16 tumors treated with ICB. We observed a significant decrease in total CD45+ lymphocytes in *Creb5*-OE tumors compared to control tumors (**Fig. 5F**), consistent with our IF data showing fewer total CD45+ lymphocytes at the center of tumor (**Fig. 5B**). Further analysis of different immune cell types revealed significant changes in the composition of myeloid cells and T cells (**Fig. 5G**) in *Creb5*-OE tumors relative to control. Among myeloid cells, we observed a significant decrease in F4/80+macrophages (**Fig. 5H**), again consistent with our IF data (**Fig. 5C**), and a significant increase in Ly6G+ myeloid-derived suppressor cells (MDSCs) (Fig. 5I), but no difference in dendritic cells (DCs) (**Fig. 5J**) in *Creb5*-OE tumors relative to control. These observations suggest that *Creb5*-OE promotes the formation of an immunosuppressive TME. Within the T cell population, we again observed a trend towards an increased frequency of CD8+ T cells (**Fig. 5K**), with only minor trends for Treg cells and Foxp3 negative CD4+ T cells (**Fig. 5L**). Overall, these observations suggest that *Creb5* overexpression disfavors anti-tumor immunity, resulting in an overall decrease in immune infiltration and an increase in potentially inhibitory populations such as MDSCs.

Collectively our data suggest that Creb5 overexpression can promote immune evasion and ICB resistance through the upregulation of collagen production in the ECM. We next sought to address the mechanism by which tumor-intrinsic collagen production inhibits anti-tumor immunity. Collagen is a ligand for the ITIM-containing inhibitory receptor LAIR1, which is broadly expressed on immune cells, including CD8^+^ T cells, NK cells, and macrophages, and inhibits their activation(*28–31*). Blocking LAIR1 signaling inhibits tumor progression(*32–37*). Immune cells increase expression of LAIR1 upon sensing collagen(*30*, *38*). Therefore, we assessed the expression of LAIR1 on immune cells in Creb5-OE and control tumors. At baseline, all immune cells express LAIR1, with highest expression on myeloid cells, followed by CD8+ T cells, then NK cells and CD4 T cells (**Fig. 5M**). Consistent with previous reports, we observed a significant increase in LAIR1 expression overall in CD45+ cells in Creb5 OE tumors relative to control tumors (**Fig. 5N**). LAIR1 expression was elevated in Creb5 OE tumors across several individual immune cell subsets (**Fig. 5O**), with the most significant increases on macrophages and CD8+ T cells, which is consistent with our immune profiling data which show the most consistent changes in these populations (**Fig. 5A-L**). The changes in myeloid cell numbers and phenotype are intriguing because previous studies have reported that LAIR1 is involved in governing macrophage polarization(*31*, *39*, *40*). In line with these studies, we found a significant decrease in F4/80 positive macrophages and a significant increase in MDSCs which are abnormally activated monocytes with immunosuppressive phenotype (**Fig. 5H-I**). Together these findings suggest that Creb5 OE in tumors promotes an immunologically cold TME, with lower numbers of effector lymphocytes and leukocytes, and suggests that these alterations in immune composition may be related to the expression of LAIR1 and direct engagement of collagen.

### Collagen sensing by LAIR1 is required for CREB5-mediated immunotherapy resistance

Given the increase in tumor-intrinsic collagen deposition and increased expression of LAIR1 on all immune cells in *Creb5* OE tumors, we sought to functionally test whether CREB5-mediated OE of collagen and collagen-stabilizing genes induces resistance to immunotherapy by engaging LAIR1 on immune cells. To determine whether engagement of LAIR1 is required for outgrowth of *Creb5*-OE and collagen OE B16 cells in ICB-treated mice, we performed an *in vivo* competition assay using *LAIR1*^−/−^ mice. We implanted an equal mix of B16-dCas9 cells expressing two control, *Creb5*, *Col16a1*, and *Col17a1* asgRNAs into *LAIR1*^−/−^ or WT mice treated with GVAX and anti-PD-1 (**Fig 6A**). We included asgRNAs targeting the transmembrane collagen COL17A1, a known ligand for LAIR1(*31*). As expected, *Creb5* asgRNA was enriched relative to control asgRNAs in ICB-treated tumors compared to NSG (**Fig. 6B** asgRNA 1 for each gene**, S6A** asgRNA 2 for each gene). However, we found the competitive advantage for *Creb5*-OE cells was reduced by 50% in *LAIR1*^−/−^ mice relative to control (**Fig. 6B, S6A**), consistent with the model that CREB5-mediated resistance partially depends on collagen sensing by LAIR1. Moreover, we found that asgRNAs targeting *Col16a1* or *Col17a1* alone were also enriched in ICB-treated mice relative to NSG mice, and that the competitive advantage for *Col16a1-* and *Col17a1*-OE cells was similarly reduced in *LAIR1*^−/−^ mice (**Fig. 6B, S6A**). Thus, immunotherapy resistance mediated by both CREB5, and collagen overexpression involves tumor-intrinsic collagen sensing by the inhibitory receptor LAIR1 (**Fig. 6B**).

**Fig 6:**
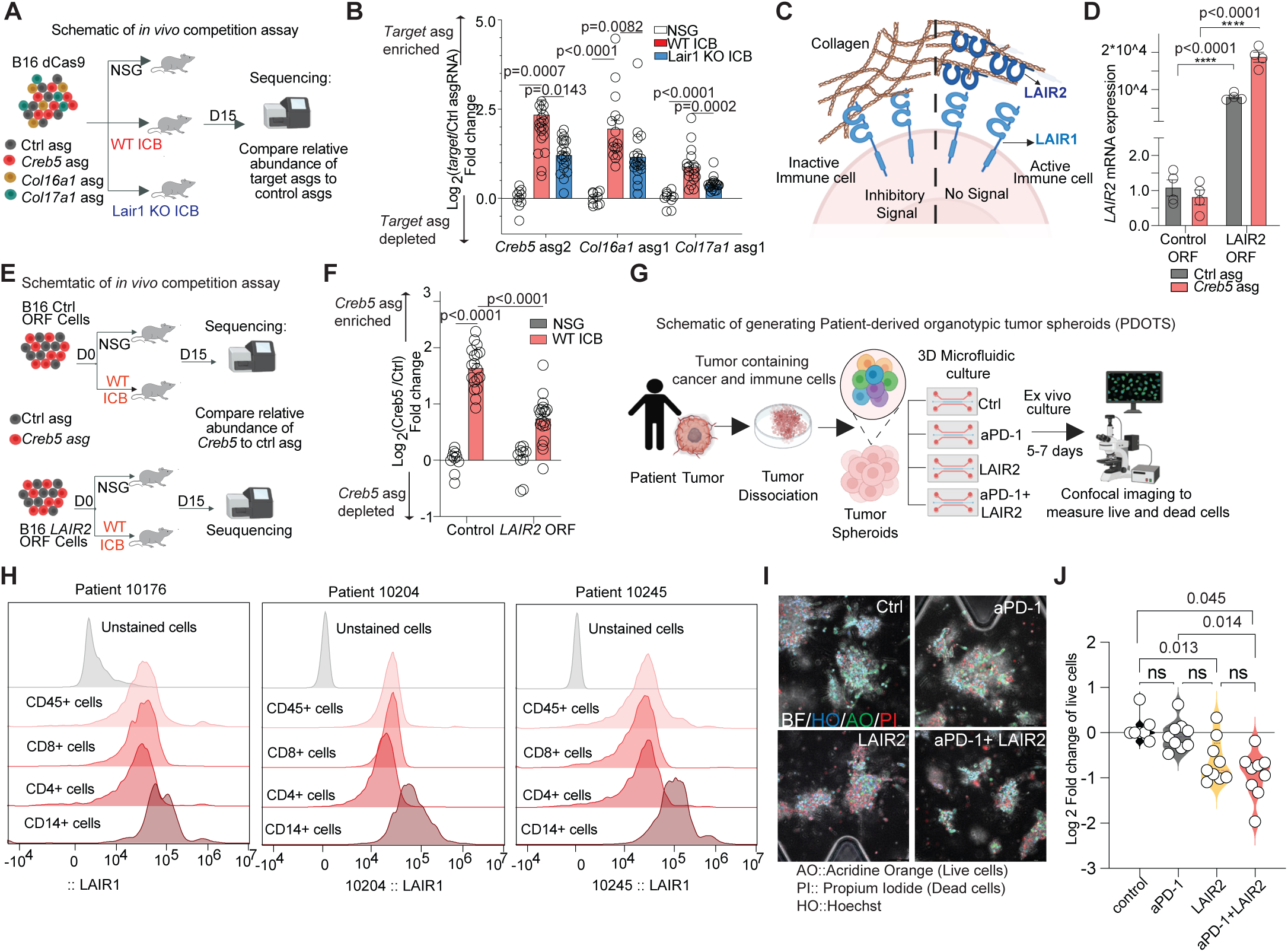
Collagen sensing by LAIR1 is required for CREB5-mediated immunotherapy resistance. A. Schematic of *in vivo* competition assay. B. Results of *in vivo* competition assay showing log2 fold change for *Creb5* asgRNA (Left), *Col16a1* asgRNA (Middle), and *Col17a1* asgRNA (Right) relative to control in B16 tumors. C. Schematic of collagen LAIR1 signaling. D. Transcript abundance for human *LAIR2* in Control (Left) or *LAIR2* ORF B16 cells. E. Schematic of *in vivo* competition assay. F. Results of *in vivo* competition assay showing log2 fold change for *Creb5* asgRNA relative to control in B16 Control ORF tumors (Left) or B16 *LAIR2* ORF tumors (Right). G. Schematic of PDOTs culture. H. Histogram plot showing the LAIR1 staining in patients tumor infiltrating lymphocytes. I. Representative IF images of PDOTs with no treatment or aPD-1 treatment (Top) or treated with recombinant human LAIR2 (2ug/ml) alone or in combination with aPD-1 (Bottom). J. Quantification of PDOTs IF images.

To further confirm the requirement of collagen sensing by LAIR1, we blocked LAIR1 signaling *in vivo* by engineering B16 cells to express LAIR2, a secreted decoy receptor that competes with LAIR1 for collagen binding (**Fig. 6C-D**)(*41*). LAIR2 is not present in mice and has a higher binding affinity for collagen than LAIR1(*41*). Thus, in engineered tumors secreting LAIR2, binding of LAIR1 to collagen will be reduced. Using these cells, we performed an in vivo competition assay with mixed *Creb5* and control asgRNA cells in both control background B16 tumors and B16 cells engineered to secrete LAIR2 (**Fig. 6E**). We observed a ∼50% reduction in the competitive advantage for *Creb5* OE cells in LAIR2 secreting tumors (**Fig. 6F**), similar to the results we obtained using *LAIR1*^−/−^ mice. These findings implicate collagen upregulation and engagement of LAIR1 as one of the major mechanisms for CREB5-mediated immunotherapy resistance.

Finally, to assess the importance of LAIR1 signaling in primary patient tumor samples, we took advantage of the patient-derived organotypic spheroid (PDOTS) microfluidic imaging platform(*42*). PDOTS are isolated from human tumors and retain autologous lymphoid and myeloid cell populations. This platform uses a three-dimensional microfluidic culture system that enables the therapeutic manipulation of primary patient tissues for the preclinical testing of therapeutics (**Fig. 6G**)(*42*). We set out to perform killing assays using PDOTS treated with anti-PD-1 or recombinant LAIR2 to block LAIR1 signaling. For these studies, we isolated PDOTS from three patients previously treated with either a MEK inhibitor or anti-PD1 therapy (**Table S14**) and confirmed the expression of *Creb5* in cancer cells (**Fig. S6B**), and LAIR1 expression on immune cells (**Fig. 6H**). We treated PDOTs with an anti-PD-1 antibody with or without recombinant LAIR2. While anti-PD-1 treatment alone did not have a significant effect on tumor killing in PDOTS, we observed significantly increased killing in conditions treated with LAIR2 alone or LAIR2 and anti-PD-1 (p=0.0023) (**Fig. 6H-I).** These findings suggest the collagen-LAIR1 signaling axis is a potential therapeutic target in ICB resistant patients.

## DISCUSSION

Here we performed an unbiased *in vivo* genome-wide CRISPR activation screen to identify novel mechanisms of ICB resistance. The transcription factor CREB5 was the overall top resistance-causing hit in our *in vivo* CRISPRa screens, and our mechanistic studies revealed that *Creb5* overexpression drives a mesenchymal-like transcriptional program characterized by high expression of collagens and collagen-stabilizing factors. Using genetically modified tumors and mice, we found that tumor-intrinsic collagen deposition is responsible for CREB5-mediated immunotherapy resistance, and that this is at least in part due to direct inhibitory signaling from the immune receptor LAIR1. The structure and collagen density of the tumor-specific ECM has been associated with poor prognosis in several types of cancer(*43–46*). However, the source of collagen has often been assumed to be stromal cells. Here we report that cancer cells can deploy resistance mechanisms such as increased expression of the transcription factor CREB5 to remodel the tumor ECM; and that collagen deposition by cancer cells is important for CREB5-mediated ICB resistance. These findings are concordant with a prior report showing that oncogenic collagen I homotrimers from cancer cells impair tumor immunity in pancreatic cancer(*47*). By manipulating collagen expression or blocking collagen deposition, we confirmed that collagen deposition is important for CREB5-mediated immune escape and is a major mechanism of ICB resistance in melanoma cells.

Collagens in the ECM provide a physical barrier to tumor immune infiltration, while also acting as a ligand for immune inhibitory receptors such as LAIR1(*31*). LAIR1 binds to collagens via the conserved Gly-Pro-Hyp collagen motif and functions as an inhibitory receptor, activating the phosphatase SHP-1 through its cytoplasmic immunotyrosine-based inhibitory motif (ITIM)(*31*, *48*). Prior studies have reported that blocking LAIR1 remodels the tumor collagen matrix, enhances tumor infiltration and activation of CD8^+^ T cells, and repolarizes suppressive macrophage populations, resulting in increased survival across murine models of colon and mammary carcinoma(*30*, *33*). Here, we demonstrate that cancer cells themselves remodel the tumor ECM through CREB5-mediated transcriptional programs that engage the collagen-LAIR1 signaling axis. In addition, blocking collagen-LAIR1 signaling in patient-derived tumor organoids promotes cancer cell death, suggesting that blocking LAIR1 is a promising potential therapeutic strategy in melanoma.

Interestingly, CREB5 has previously been implicated in cancer progression and therapy resistance. Recent studies revealed frequent CREB5 amplification and overexpression in kidney cancers, sarcomas, lymphomas, lung adenocarcinomas, glioblastomas, and gliomas(*49*). Other studies have reported upregulation of CREB5 expression in ovarian cancers(*50*), hepatocellular carcinoma(*51*), and colorectal cancer(*49*) and its role in promoting cancer cell proliferation and metastasis. Moreover, we previously identified the role of CREB5 in prostate cancer in promoting resistance to androgen receptor-targeting therapies(*11*). Together, these studies implicate CREB5 in therapy resistance in advanced cancers of several lineages. Whether the CREB5-driven mesenchymal-like phenotype and collagen deposition we describe here also participate in advanced disease in other cancer types is currently unknown. We observed that CREB5 overexpression caused upregulation of several pathways including EMT, MYC, and NFκB. Thus, CREB5 may promote resistance by several mechanisms depending on the cell lineage and context. Further studies will be required to examine the role of CREB5 in inducing ICB resistance in other cancer types.

The identification of a transcription factor as the top overall resistance-causing hit highlights the complexity of cancer immune evasion and the importance of functional studies to determine the causal nature of transcriptional changes associated with resistance. Indeed, while some previous studies have been able to identify single gene loss of function or gain of function alterations associated with ICB resistance(*52–54*), several other large studies suggest that such single gene resistance mechanisms to ICB are rare(*8*, *55–58*). Here using gain-of-function screens, we functionally linked a transcription factor driven mesenchymal and collagen-high transcriptional state to ICB resistance and discovered key mediators of that state that may be targeted with therapeutics, such as LOXL1(*59*) and LAIR1(*33*).

## Acknowledgements

The authors would like to thank all members of the Manguso Lab and Hahn Lab at MGH, DFCI, and the Broad Institute. P. Rogers and the staff of the Broad Flow Cytometry core for assistance with cell sorting; T. Caron and the staff of the Broad Institute Comparative Medicine programme for assistance with animal studies. Graphics in Figs. 1a, 1d, 4d, 5d, 5h were created using BioRender (https://www.biorender.com).

## Contributions

Study design: PT, WCH, and RTM

Design of in vivo screens: PT, GD, WCH, RTM

IHC/IF imaging - PT, JMC, NALF, EK

CUT&RUN PCR - SMM

Execution of in vitro and in vivo studies: PT, KJC, NALF, CLC, SH, ZS, KBY, SA, JT

Patient-derived organotypic tumor spheroids planning and execution: RWJ, AMC, GMB, CAP, OYR

Computational analysis: SK, SSF, MSF, NH, AVK, NALF

LAIR1 knockout mice: PT, KHC, CLC, MV, SD

Manuscript writing and revision: PT, JT, KJC, KBY, WCH and RTM

## Data and materials availability

All data and materials generated in this study are available upon request from the corresponding authors. Raw data from Supplementary Tables S1, S3, S5, and S7 have been deposited to GEO accession XX.

## Supplementary Materials

Materials and Methods

Figs. S1 to S5

Tables S1 to S14

## Materials and Methods

### Cell lines

The B16-F10 (B16) melanoma and B16 granulocyte–macrophage colony-stimulating factor-secreting cells (GVAX) were a gift from S. Dougan and G. Dranoff (Novartis Institutes for Biomedical Research). B16 cells were previously subcloned to minimize screening cell population heterogeneity (https://www.nature.com/articles/s41590-022-01315-x#citeas). The YUMMER1.7 Braf/Pten melanoma cell line (YUMMER) was a gift from M. Bosenberg (Yale University) and S. Kaech (Salk Institute for Biological Studies). A375 melanoma cells were purchased from ATCC. All cell lines were tested for mycoplasma and cultured in DMEM (Gibco), 10% fetal bovine serum, and antibiotics.

### Mice

All *in vivo* studies conducted at the Broad Institute were approved by the Broad Institute IACUC committee and mice were housed in a specific-pathogen free facility. All *in vivo* studies at MGH were conducted according to protocols approved by the MGH Institutional Animal Care and Use Committee. Mice were housed in a specific pathogen-free facility with 12-h dark/12-h light cycles and optimal ambient temperature (21 °C ± 1 °C) and humidity (40% ± 10%) conditions. Seven- to 12-week-old WT C57BL/6J mice were obtained from Jackson Laboratories. Colonies of NOD.Cg-Prkdcscid Il2rgtm1Wjl/SzJ (NSG) immunodeficient mice and Igs2tm1.1(CAG-cas9*)Mmw/J mice were bred at the Broad Institute. For experiments at MGH, NSG mice were obtained from Jackson Laboratories. LAIR1 KO mice were a generous gift from S. Demehri and were bred at MGH (doi: 10.1161/ATVBAHA.117.309253). For tumor challenges with WT or LAIR1 KO mice, age-matched female mice were used. For tumor challenges with YUMMER cells, age-matched male Cas9 mice were used. Both male and female age-matched mice were used for NSG tumor challenges.

### Tumor Challenges and Treatment

For screening experiments, 2.0 X 10^6^ library-transduced B16-dCas9 cells were resuspended in a 50:50 mix of phosphate-buffered saline (PBS) and growth factor-reduced Matrigel (Corning). Tumor cells were subcutaneously injected into the flank (bilateral) and measured manually every 3-4 days (beginning 6 days post-tumor challenge) until survival end-point was reached or no observable tumor remained. Tumor volumes were estimated using longest dimension (length) and the longest perpendicular dimension (width), using the formula (L × W^2^)/2. Tumor volumes were assessed every 3–4 days until either the survival end point was reached, or no palpable tumor remained. On days 1 and 4, treated mice received subcutaneous injection in the abdomen of 2.0 × 10^5^ GVAX cells that had received 35 Gy of irradiation. These mice were also administered 200 μg of rat monoclonal anti-PD-1 (Bio X Cell, clone 29F.1A12) on days 6, 9, and 12 via intraperitoneal (i.p.) injection.

Tumors were collected 12–15 days after implantation, minced with scissors and pooled by equal tissue mass in groups of 5–10 tumors within each experimental arm. Pooled tissue was then digested with Proteinase K (Qiagen) and buffer ATL (Qiagen), and genomic DNA was extracted using the Qiagen Blood Maxi kit. From 40 to 240 μg of genomic DNA per pooled sample, the sgRNA region was PCR amplified (using the P5 and P7 Illumina primers; sequences are available at https://portals.broadinstitute.org/gpp/public/resources/protocols) and sequenced using an Illumina HiSeq.

For *in vivo* competition experiments, 2.0 X 10^6^ tumor cells were resuspended in PBS and injected subcutaneously into both sides of the flank. For experiments with B16 cells, the above outlined treatment regimens were carried out. For all other cell lines, treated mice received i.p. injections of anti-PD-1 on days 6, 9, and 12 post tumor implantation. For competition studies with CD8-depletion arm, mice were treated with 200 μg of anti-CD8b (Bio X Cell, clone 53-5.8), via i.p. injection beginning 1 day prior to tumor challenge, continuing every fourth day for the duration of the experiment.

### Genomic DNA isolation from tumors

Tumors and in vitro cultures were digested overnight in a 1:10 solution of Proteinase K (Qiagen) and buffer ATL (Qiagen). Genomic DNA was extracted from digested samples using DNEasy Blood and Tissue genomic DNA isolation kits (Qiagen).

### In vivo CRISPR screening

For the primary genome-scale *in vivo* screen, a library of sgRNAs targeting 18,000 genes with 4 sgRNAs per gene along with both non-targeting and intergenic-targeting control guides was cloned into the pXPR_502_sgRNA vector. This library was separated into four pools that had 22,313, 22,277, 22,251 and 22,227 unique sgRNAs represented in them (GPP Caprano SetA/B).

For the secondary subgenome-scale *in vivo* screen, we selected 1). 200 most depleted and 200 most enriched genes from STARS analysis of primary genome-scale CRSIPRa screen, 2). 200 most depleted and 200 most enriched genes from hypergeometric analysis of primary CRISPRa screen, 3). 50 most depleted hits from each individual pool of primary CRISPRa screen, 4). 50 most enriched hits from each individual pool of primary CRISPRa screen, 5). 100 most depleted genes from STARS analysis of each of the CRISPR KO screens from 8 cancer cell lines, 6). 50 most enriched genes from STARS analysis of CRISPR KO screen in each of the 8 cell lines. After deleting the genes duplicated in all these settings, we got a final list of 2,118 genes to which we added control guides. Together, we generated a library of 21,323 asgRNAs targeting 2,118 genes with 10 sgRNAs per gene along with control guides cloned into the pXPR_502_sgRNA vector (GPP). These pools were delivered to B16 cells that stably expressed dCas9. Transduced cells were selected in blasticidin and puromycin and expanded *in vitro* for 7-10 days before *in vivo* transplantation into mice. Cells were maintained in culture at >2,000× coverage per sgRNA for the duration of the screen.

### Analysis of screening data

Guide sequences were demultiplexed and quantified using PoolQ2 (https://portals.broadinstitute.org/gpp/public/software/poolq). Read counts were library normalized per 1 million reads and log2 transformed with a pseudocount of one. Gene-targeting guides were z normalized by the control sgRNA distribution. Guide fold changes were calculated as residuals fit to a natural cubic spline with 4 degrees of freedom. For the genome-scale primary screen, fold changes from all four pools were quantile normalized with gene-wise mean imputation before downstream analyses. Enriched and depleted genes for each screen were calculated using the STARS algorithm (GPP).

### Generation of overexpression vectors

For the overexpression vectors pLX_313_mCD19_truncated and pLX_313_Lair2, cDNA (Origene) was PCR amplified using primers containing attB sites and cloned into the pLX_313-Gateway destination vector using Invitrogen BP Clonase II and LR Clonase II Gateway reactions according to the manufacturer’s instructions.

### Generation and validation of CRISPR-a and CRISPR-KO cell lines

For CRISPR activation experiments, B16.1, B16F10, YUMMER, and A375 cells were sequentially lentivirally transduced with pXPR_109_dcas9 (GPP) and pXPR_502_sgRNA (GPP) with polybrene and selected in blasticidin and puromycin. For LAIR2 validation experiments, B16.1 CRISPRa cells with lentivirally transduced with pLX_313_LAIR2 or pLX_313_mCD19_truncated and selected in hygromycin. For LOXL1 KO experiments, B16.1 cells were transiently transfected with Cas9 and sgRNA vector (pSpCas9(BB)-2A-Puro (PX459) targeting Loxl1 or control with Lipofectamine transfection reagent (Thermo Fisher Scientific, L3000015). Transfected populations were selected in antibiotics for 2–4 d and bulk transfectant populations were used for subsequent experiments. For experiments with LOXL1 ko cells overexpressing CREB5, control and LOXL1 KO cells were lentivirally transduced with pXPR_109_dcas9 (GPP) followed by pXPR_502_sgRNA targeting *Creb5* or control. For *Creb5* CRISPR KO studies, B16.1, YUMMER, and A375 cells were lentivirally transduced with pSCAR_Cas9_BSD (Addgene) and pXPR_BRD060 (Addgene) targeting control or *Creb5*. Prior to in vivo experiments, CRISPR KO cells were lentivirially transduced with IDLV-Cre (Addgene). All cell lines were validated for overexpression and/or knockout efficiency via RT–qPCR, flow cytometry, or western blot.

### CRISPR sgRNA sequences

*NT sgRNA1 : 5’-TTGTTACGATATTCGCGCGA-3’*

*NT sgRNA2 : 5’-ATATCGTATCGACTCGATTC-3’*

*NT sgRNA3 : 5’-AAACGAGGCTGTTCGTACAC-3’*

*Old Control sgRNA2 : 5’-AAACTCATACGTAGCGAATC-3’*

*IT sgRNA1 : 5’-TCAGACCTACTTGTGATGGG-3’*

*IT sgRNA2 : 5’-GGCAGCAGGAGGGGCAACAA-3’*

*IT sgRNA3 : 5’-AGGAGGTGGCGCGCTCTTGG-3’*

*Creb5 sgRNA1 : 5’-GCTCTGGCTCTCTCTCCCTC-3’*

*Creb5 sgRNA2 : 5’-AGAGTGAGCAGCAGAGCCAG-3’*

*Col16a1 sgRNA1 : 5’-CCGCAGCCGCCGAACCCTCG-3’*

*Col16a1 sgRNA3 : 5’-AGCGCGGCGGTGCAGGCCAA-3’*

*Col17a1 sgRNA1 : 5’-AATGGTTAATCTACCTGAGC-3’*

*Col17a1 sgRNA4 : 5’-AGGAACCACACTTTATAGAC-3’*

*Col4a1 sgRNA2 : 5’-GTCGCAGGGAGTCCGAACGC-3’*

*Col4a1 sgRNA4 : 5’-CTGCGACACCAAGTCCCGAG-3’*

*Col1a1 sgRNA2 : 5’-GGAACCCTGCCTCTCGGAGA-3’*

*Control KO sgRNA1 : 5’-CGAGGTATTCGGCTCCGCGG-3’*

*Control KO sgRNA5 : 5’-ATTGTTCGACCGTCTACGGG-3’*

*Loxl1 KO sgRNA2 : 5’-CGTCACGGACCTACGAACA-3’*

*Creb5 KO sgRNA : 5’-AATGCTTACAGGCATCAAGA-3’*

*CONTROL KO sgRNA1 : 5’-CGGGCAGAACGACCCTGACG-3’*

*CONTROL KO sgRNA2 : 5’-CGCGCACCACGGGCGCGCAC-3’*

*CREB5 KO sgRNA1 : 5’-AATGGGAACATGAACACCAT-3’*

*CREB5 sgRNA4 : 5’-CACTGAGCTAAAAAGGAGAA-3’*

*CONTROL sgRNA1 : 5’-AAAACAGGACGATGTGCGGC-3’*

### RNA Seq Sample Preparation and Analysis

B16.1 cells transfected with *Creb5* or control asgRNA were seeded in 6-well plates and cultured for 24-48 hours. RNA was extracted from cell pellets using the RNeasy plus mini kit (Qiagen, 74034) according to the manufacturer’s instructions. First-strand Illumina-barcoded libraries were generated using the NEB RNA Ultra II Directional kit according to the manufacturer’s instructions using a 10-cycle PCR enrichment. Libraries were sequenced on an Illumina NextSeq 500 instrument using paired-end 37-base pair reads. Read data were adapter and quality trimmed using Trimmomatic (v.0.36). Trimmed reads were quantified by pseudoalignment to mm10 using Kallisto (v.0.46.0). RNA-seq counts were quantified using the program tximport. A variance stabilizing transformation was applied to the counts matrix, and differentially expressed genes were quantified using the DESeq2 R package. Gene set enrichment analysis was performed using GSEAPreranked. The metric of one minus Pearson correlation between samples was used to perform hierarchical clustering of the RNA-seq samples.

### Analysis of patient bulk RNA-seq

Expression values (TPM) and clinical annotations were downloaded from (Liu et al, 2019). Patients were divided into CREB5 high and CREB5 low subsets based on the CREB5 median expression value. Progression-free survival (PFS) was compared between the low and high CREB5 expression groups. The Python3 package, lifelines, was used to fit a Kaplan-Meier estimator and to calculate p-value via log-rank test. The relationship between CREB5 expression and PFS was also measured by a multivariate regression using the Cox proportional-hazards model (from the lifelines package) with the additional variables of tumor purity and metastatic stage as presented in the clinical annotations. Additionally, a Mann-Whitney U test from the Python3 package scipy.stats was used to assess differences in CREB5 expression between responders and progressors to immunotherapy. Responders were classified as those with partial response (PR) or complete response (CR) and progressors were those with progressed disease (PD). GSEA was also used to identify genesets which correlated with PFS. For all genes in the geneset, a multivariable CoxPH model was fitted on that gene’s expression as well as tumor purity and metastatic stage. Genes were ranked using the output from the CoxPH model using the equation log2(p-val) * sign(log (HR)). GSEA was run on all the Hallmark genesets as well as a set of melanoma specific genesets from Pozniak et al, 2022. Genesets with positive normalized enrichment scores are associated with positive hazard and worse PFS.

### Analysis of patient-derived scRNA-seq

Cell annotations and associated expression values (TPM+1 normalized) were downloaded from GSE115978 as well as from Pozniak et al, 2022. Dataset was filtered to cells annotated as malignant cells. Using the following procedure, gene set enrichment analysis was performed to assess the correlation of CREB5 expression with the MSigDB Hallmark genesets and genesets from (Pozniak et al, 2022). All cells with CREB5 TPM = 0 were removed. Only tumors with 10+ malignant cells expressing CREB5 were considered. For each tumor, the correlation between CREB5 and each other gene was calculated using the Pearson correlation. The correlation coefficient for each gene was averaged among all tumors. A ranked gene list was created and sorted by the average CREB5 correlation coefficient among all tumors. GSEA preranked was run on this sorted list. For the GSE115978 dataset, the correlation of all transcription factors with the Mesenchymal signature was performed using the same procedure. A list of transcription factors was drawn from (Lambert et al, 2018). TFs with five or less samples with 10+ malignant cells were excluded from this analysis.

### RNA isolation and RT–qPCR

For RT-qPCR experiments, cells were seeded in 6-well plates and cultured for 24-48 hours. Total RNA was isolated from the cells using the RNeasy plus mini kit (Qiagen, 74034) per the manufacturer instructions. PCR reactions were performed using Power SYBR Green PCR Master Mix (Applied Biosystems, 4367659) and detected on a QuantStudio 6 Flex Real-Time PCR System (Applied Biosciences, 4485691). Gene expression levels were normalized to *18s*. The primers used to detect mRNA expression are listed below.

### RT–qPCR Primers

*18s*-Forward: 5’-GTAACCCGTTGAACCCCATT-3’

*18s*-Reverse: 5’-CCATCCAATCGGTAGTAGCG-3’

*Col1a1*-Forward: 5’-GTGCGATGACGTGATCTGTGA-3’

*Col1a1*-Reverse: 5’-CGGTGGTTTCTTGGTCGGT-3’

*Loxl1*-Forward: 5’-CCACTACGACCTACTGGATGC-3’

*Loxl1*-Reverse: 5’-GTTGCCGAAGTCACAGGTG-3’

*Serpine1*-Forward: 5’-ACCGCAACGTGGTTTTCTCA-3’

*Serpine1*-Reverse: 5’-TTGAATCCCATAGCTGCTTGAAT-3’

*Col4a1*-Forward: 5’-GGGATGCTGTTGAAAGGTGAA-3’

*Col4a1*-Reverse: 5’-GGTGGTCCGGTAAATCCTGG-3’

*Creb5*-Forward: 5’-GTCCCAGGCTCTCTATCATCTC-3’

*Creb5*-Reverse: 5’-ATAGGCATCAAGACGGCAGAA-3’

*Col16a1*-Forward: 5’-GAGAGCGAGGATACACTGGC-3’

*Col16a1*-Reverse: 5’-CTGGCCTTGAAATCCCTGG-3’

*Creb5*(ORF)-Forward: 5’-TTCTGCCGTCTTGATGCCTA-3’

*Creb5*(ORF)-Reverse: 5’-ATCATGTGTCCGATGGTGCT-3’

*18S*-Forward: 5’-GTAACCCGTTGAACCCCATT-3’

*18S*-Reverse: 5’-CCATCCAATCGGTAGTAGCG-3’

*COL1A1*-Forward: 5’-GTGCGATGACGTGATCTGTGA-3’

*COL1A1*-Reverse: 5’-CGGTGGTTTCTTGGTCGGT-3’

*COL4A1*-Forward: 5’-GGGATGCTGTTGAAAGGTGAA-3’

*COL4A1*-Reverse: 5’-GGTGGTCCGGTAAATCCTGG-3’

*LOXL1*-Forward: 5’-CCACTACGACCTACTGGATGC-3’

*LOXL1*-Reverse: 5’-GTTGCCGAAGTCACAGGTG-3’

*SERPINE1*-Forward: 5’-ACCGCAACGTGGTTTTCTCA-3’

SERPINE1-Reverse: 5’-TTGAATCCCATAGCTGCTTGAAT-3’

*CREB5*-Forward: 5’-TGGGACACATGATGGAGATGA-3’

*CREB5*-Reverse: 5’-GGTGTGCATGAAGGTGGGAA-3’

*LOXL2*-Forward: 5’-GGGTGGAGGTGTACTATGATGG-3’

*LOXL2*-Reverse: 5’-CTTGCCGTAGGAGGAGCTG-3’

*LAIR2*-Forward: 5’-ATGTCTCCACACCTCACTGCTC-3’

*LAIR2*-Reverse: 5’-TGGTGCATCAAATCCGGAGGC-3’

### Western blot

Whole cell lysates from tumor cells were prepared in RIPA Lysis and Extraction Buffer (ThermoFisher Scientific), Halt™ Protease Inhibitor Cocktail (ThermoFisher Scientific), and PhosSTOP™ (Roche), and EDTA (xxx). Protein concentration was measured using a BCA protein assay kit (Pierce) for the addition of NuPAGE™ LDS Sample Buffer (ThermoFisher Scientific) and Dithiothreitol (DTT). Samples were heated at 70℃ for 10 minutes and 50-100 μg of protein was loaded and separated on either 4%–12% Bis-Tris Plus Gels for low molecular weight proteins or 3%–8% Tris-Acetate Gels for high molecular weight proteins (Life Technologies) in NuPAGE™ MOPS SDS running buffer (Life Technologies). Protein was transferred to 0.45-mm nitrocellulose membrane (Bio-Rad) using the iBlot 2 (ThermoFisher Scientific) for low molecular weight proteins or via wet transfer (10V for 4 hrs) for high molecular weight proteins. Membranes were washed with Tris-Buffered Saline plus 0.1% Tween 20 (TBST) and blocked in Odyssey Blocking Buffer (LI-COR) for 30 min. Membranes were incubated in primary antibodies overnight at 4℃ and subsequently washed and incubated in secondary antibodies. Blots were visualized using the Odyssey CLx scanner (LI-COR) and analyzed using Image Studio Lite. The following primary antibodies were used for detecting designated protein expression in western blot assays: CREB5 (Novus Biologicals, H00009586-M02, Abnova), CREB5 (ThermoFisher, PA-565593, Invitrogen), Anti-Mouse GapDH (Abcam, ab8245), Anti-Rabbit GapDH (Abcam, ab9485), COL1A1 (ThermoFisher, PA5-29569), COL4A1 (ThermoFisher, PA5-104508), COL16A1 (ThermoFisher, PA5-103382). For flow cytometry, the following anti-mouse primary antibodies were used: LAIR1 (clone 113, ThermoFisher).

### Tumor Immunofluorescence staining

Whole B16.1 tumors were fixed overnight in 10% neutral-buffered formalin and then transferred to 70% ethanol until processing. Fixed tissue was subsequently embedded into paraffin, sectioned, and then mounted onto slides for staining against mouse COL1A1 (Cell Signaling Technology, 72026S), COL4A1 (Rockland, 600-401-106-0.1), PECAM1 (CD31) (R&D Systems, AF3628), FOXP3 (Cell Signaling Technology, 12653S), CD4 (Invitrogen, 14-9766-82), PTPRC (CD45) (Cell Signaling Technology, 70257S), ADGRE1 (F4/80) (Cell Signaling Technology, 70076S), CD8A (Invitrogen, 14-0808-82), and 4′,6-diamidino-2-phenylindole (DAPI) (Thermofisher Scientific, 62248). Slides were deparaffinized as follows: Heated to 60C for 15min, transferred to xylene for 270sec, then transferred to three ethanol solutions for 90sec each at 100%, 95%, and 70% concentrations respectively. Subsequently, slides were heated in a steam bath for 50min in 1:100 solution of Antigen Unmasking Solution (Vector Laboratories, H-3300-250). Due to high melanin expression in melanoma tumors, sections were then depigmented for 5 hours by immersing in 3% hydrogen peroxide solution and exposed to bright light under a LED floodlight. Slides were imaged on the Lunaphore COMET system according to manufacturer’s instructions.

Antibody dilutions were as follows: 1:100 for COL1A1, CD8A, and FOXP3, 1:200 for COL4A1, CD45, F4/80, and PECAM1, 1:1000 for DAPI, and 1:2000 for secondary antibodies Goat anti-Rabbit IgG Alexa Fluor 647+ (Invitrogen, A32733), Donkey anti-Goat IgG Alexa Fluor 647 (Invitrogen, A21447), Donkey anti-Rat IgG Alexa Fluor 647 (Invitrogen, A48272TR), and Donkey anti-Rabbit IgG Alexa Fluor 555+ (Invitrogen, A32794). Images were quantified using QuPath software version 0.5.1. Briefly, tumor regions were annotated along collagen-thick tumor edges to section inner and outer regions of the tumor. For collagen quantification, collagen signal for both COL1A1 and COL4A1 was segmented by thresholding and collagen-positive regions were classified as either endothelial-associated collagen or tumor-associated collagen based on the presence of overlapping signal with PECAM1. Cell numbers were determined using the StarDist plugin version 0.5.0. For the tumor microenvironment, an object classifier was trained using the Random Trees method to classify cells based on marker expression. F4/80 positive area was used instead of a classifier to quantify macrophage presence due to their non-circular shape.

### Tumor trichrome staining

Whole B16.1 tumors were mounted onto slides and deparaffinized as previously described and then stained with Trichrome Stain Kit (Abcam, ab150686), according to manufacturer’s instructions. Slides were imaged using the Leica Aperio Versa 8 inverted microscope.

### Flow Cytometry

For flow cytometry of B16.1 cells, trypsinized cells were washed in PBS + 2% FBS + 5 mM EDTA. Cells were stained with antibodies against surface proteins for 15 min at 4 °C per the manufacturer’s instructions. All samples were acquired using a Beckman Coulter Cytoflex instrument and analyzed with FlowJo software.

### Flow cytometry analysis of Tumor infiltrating lymphocytes

Mice were subcutaneously injected with 2 × 10^6^ B16 cells and treated with anti-PD-1 or no treatment, as above. Tumours were collected on day 15, weighed and chopped before chemical and mechanical digestion with a Tumour Dissociation kit (Miltenyi) and a gentleMACS Dissociator (Miltenyi) using the m-TDK-1 programme. The resulting cell suspension was passed through a 70-µm filter. Isolation of immune cells was performed by density centrifugation with lympholyte reagent (Cedarlane Labs). Cells were blocked with TruStain FcX Plus (anti-mouse CD16/32) antibody (1:100, BioLegend) in PBS with 2% FBS. Dead cells were excluded using Live/Dead Fixable Blue Dead Cell stain (1:1,000, ThermoFisher) added concurrently with surface antibodies. Samples were stained with indicated antibodies for 30 min on ice. After washing, cells were fixed using a FOXP3/Transcription Factor Staining buffer set (eBiosciences) as per the manufacturer’s instructions, blocked with TruStain FcX, mouse and rat serum, then stained with intracellular antibodies before proceeding with blocking and staining for intracellular and extracellular markers. Analysis was done on a Beckman Coulter Cytoflex LX flow cytometry system using single-colour compensation controls and fluorescence-minus-one thresholds to set gate margins. One-way analysis of variance was used to make comparisons between groups. Generated *P* values were adjusted for multiple comparisons.

The following primary antibodies were used to detect designated protein expression, anti-mouse fluorochrome-conjugated antibodies were used: CD8α (clone 53-6.7, BioLegend or BD Biosciences); CD4 (clone RM4-5 or GK1.5, BioLegend); TCRβ (clone H57-597, BioLegend); CD45 (clone 104, BioLegend); NK1.1 (clone 108741, BioLegend); FOXP3 (clone JFK-16s, eBioscience); CD3 (clone 17A2, BioLegend); IFNγ (clone XMG1.2, BioLegend); and anti-rabbit IgG (H+L), F(ab′)2 fragment (Cell Signaling, 4412).

### Patient-derived organotypic tumor spheroid (PDOTS) preparation and microfluidic device culture

PDOTS were generated as previously described (Jenkins *et al*. 2018). In brief, fresh tumor specimens from mice or humans were received in full DMEM media (Corning, 10-013-CV), supplemented with 10% BenchMark™ Fetal Bovine Serum (GeminiBio, 100-106) and 1% penicillin-streptomycin (Thermo Scientific, 15140122) on ice and minced in a 10 cm plate using sterile forceps and scalpels, if samples arrived late, they were stored in tissue storage media (Miltenyi, 130-100-008) and processed the day after. Minced tumors were resuspended in full DMEM and were passed over 100-μm and 40-μm filters sequentially to generate the S1 (>100 μm), S2 (40-100 μm), and S3 (<40 μm) fractions. S2 and S3 were washed off the filter with fresh full media and were rested in ultra-low attachment plates in (Corning, 3471) in the incubator until loading into the device. S1 fraction was washed from the filter using fresh full media supplemented with 100 U/mL collagenase type IV (Life Technologies, 17104019), and 15 mM HEPES (Gibco, 1560-080). S1 with collagenase was incubated for 15-30 minutes at 37°C followed by the addition of an equal volume of media and subsequent filtering. The S2 fraction was pelleted and resuspended in type 1 rat tail collagen (Corning, 354236) at a concentration of 2.5 or 3 mg/mL with 10x PBS + phenol red (Sigma-Aldrich, 114537-5g). A pH of 7.0-7.5 was confirmed with PANPEHA Whatman paper (Sigma-Aldrich, 2629990) after titrating the solution with NaOH (Sigma-Aldrich, 1.09138.1000). The spheroid-collagen mixture was added into the center channel of the AIM 3D microfluidic device (DAX-01, AIM Biotech). Collagen hydrogels with PDOTS were incubated for 20 minutes at 37°C in sterile humidity chambers before being hydrated with media in the side channels with or without treatments. PDOTS were treated with anti-PD-1 (250 μg/mL, pembrolizumab), 2-4 ug/ml human recombinant LAIR2 (R&D Systems, 2665-LR-050), combination of the two or left untreated. S1 and S3 were banked using BAMBANKER freezing media (Bulldog Bio, BB02) and kept for downstream analysis by flow cytometry (see relevant section).

### PDOTS viability assessment

PDOTS staining and viability analysis was done as previously described (Jenkins *et al*. 2018). In brief, staining was done in microfluidic devices by adding acridine orange/ propidium iodide (AO/PI) solution (Nexcelom, CS2-0106), diluted 1:1 in full media with 12ug/ml Hoechst (Invitrogen, H3570). Alternatively, devices were stained using 12 μg/mL Hoechst and 0.8 μg/mL PI (Thermo Scientific, P3566). After incubation with fluorescent dyes (30 minutes, 37°C), images were acquired using a Nikon Eclipse NiE fluorescence microscope, using X4 lens in 2 or 3 colors. Images were analyzed using the NIS-Elements AR software package and live/dead cell quantification was obtained by measuring the total cell area of acridine orange for live cells, PI for dead cells and Hoechst for total cells. Live cells were defined as the total area of the acridine orange channel, or the total area of Hoechst (total cells)-PI (dead cells). Percent change and log2FC (L2FC) data were generated using raw fluorescence data (live) for given treatments relative to control conditions.

### Analysis of human tumors and human T cells by flow cytometry

Single cell suspension from PDOTS processing (S3; see PDOTS generation section) was processed for flow cytometry. Single cell suspension was thawed into ImmunoCult-XF media (Stemcell Technologies, 10981) and incubated at 37°C overnight in ultra-low attachment plates (Corning, 3471) to allow cell recovery. Tumors were stained with conjugated fluorescent anti-human monoclonal antibodies against CD3 (BD Biosciences, 558117), CD4 (BD Biosciences, 557695), CD45 (BioLegend, 368513), LAIR1 (BioLegend, 342801), CD8⍺ (BioLegend, 301045), CD14+ (BioLegened, 301817). Human TILs were stained with CD3, CD4, CD45, LAIR1, CD8⍺. Samples were acquired on a Cytek Northern Lights instrument and MA900 Sony Cell sorter and analyzed with FlowJo software version 10.8.1.

### Flow cytometry and cell sorting

For flow cytometry Human TrueStain FcX (Biolegend, 422302) was used for blocking Fc receptors before labeling cells. To discriminate between live and dead cells, we used Zombie Violet Dye (Biolegend, 423114) for 15 min at 4°C, followed by surface labeling of cells for 30 min at 4°C, using standard protocols.

### RNA Isolation and RT-qPCR

Total RNA was extracted from tumor patient derived cell lines cells using the RNeasy Kit (Qiagen, 74104) according to the manufacturer’s protocol. RNA levels were quantified using the Blaze Taq one-step SYBR Green RT-qPCR kit (GeneCopoeia, QP070) according to the manufacturer’s protocol and acquired on the ROCHE Lightcycler-96 system. RNA samples were normalized to TUBB and the 2-ΔΔCt was calculated to obtain the relative gene expression.

### RT–qPCR Primers

*TUBULIN*-Forward: 5’- AAGATCCGAGAAGAATACCCTGA-3’

*TUBULIN*-Reverse: 5’- CTACCAACTGATGGACGGAGA-3’

*CREB5*-Forward: 5’- AGATGGTCCTCTGTTGGGAA-3’

*CREB5*-Reverse: 5’- TGGACACGGTTATGAGAATGA-3’

---

**Fig. S1.**
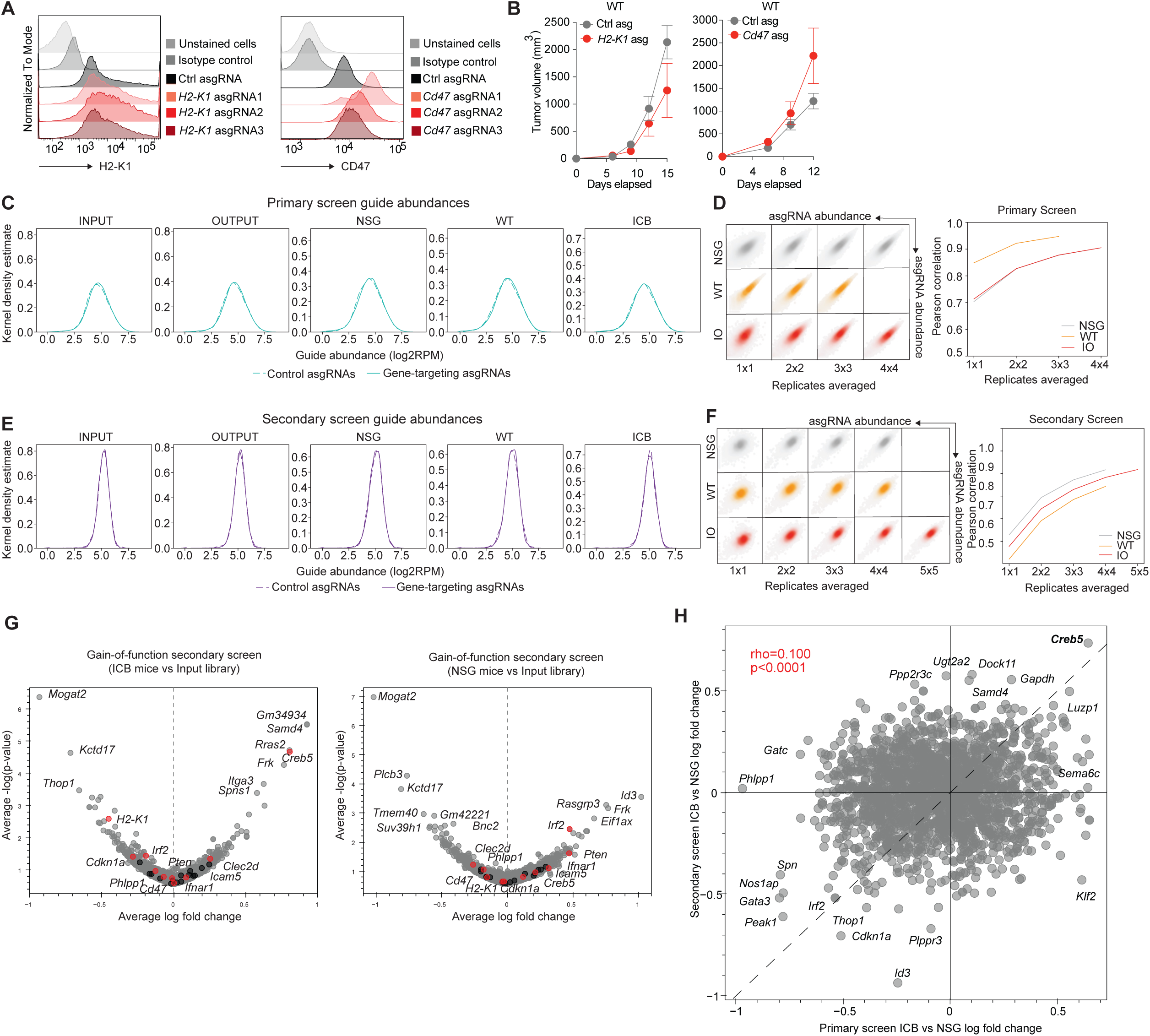
*In vivo* CRISPR activation screen identifies overexpression of Creb5 as a novel mechanism of ICB resistance. A. Histograms showing H2-K1 and CD47 staining in B16 cells. B. Tumor growth over time for untreated mice for *H2-K1* asgRNA tumors (red) relative to control (gray) (on left) and *Cd47* asgRNA tumors (red) relative to control (gray) (on right). C. Library recovery across conditions in the primary screen. Dotted lines represent control asgRNAs and solid lines represent gene-targeting asgRNAs. D. Replicate autocorrelation analysis in primary screen. Pearson correlations are calculated for the library distribution in one 8-tumor replicate versus any other replicate, two averaged replicates versus any other two, and so on (left). The mean of all possible combinations is plotted (right). E. Library recovery across conditions in the secondary screen. F. Replicate autocorrelation analysis in secondary screen (left). Pearson correlations are calculated for the library distribution in one 8-tumor replicate versus any other replicate, two averaged replicates versus any other two, and so on (left). Mean of all possible replicate combinations in the secondary screen (right). G. Volcano plot of secondary CRISPR activation screen hits in ICB mice vs. Input library (Left) and NSG mice vs. Input library (Right). Genes marked in color are known immune regulators or novel hits of interest from ICB vs. NSG library. H. Scatter plot of log fold changes of genes shared between the primary screen and secondary screen. The data have a Pearson correlation coefficient of 0.100 with p-value<0.0001.

**Fig. S2.**
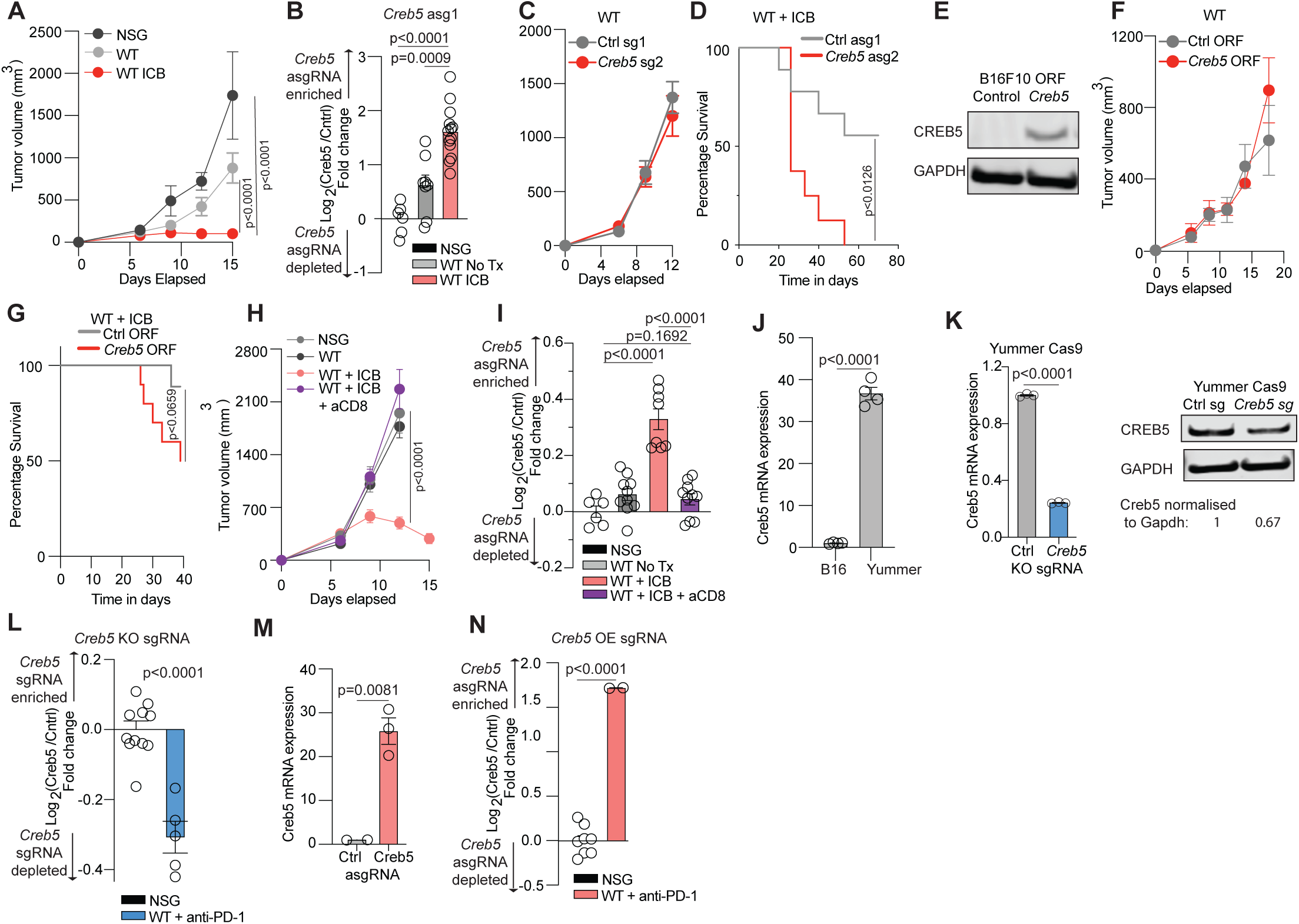
CREB5 promotes ICB resistance in melanoma cells. A. B16 Tumor growth over time in NSG (Black), WT (Grey), WT + ICB (Red) mice. B. Results of in vivo competition assay showing log2 fold change for *Creb5* asgRNA relative to control in B16 tumors. C. Tumor growth over time of *Creb5* asgRNA tumors (Red) relative to control (Gray) in untreated mice. D. Survival curve for *Creb5* asgRNA tumors (Red) relative to control (Gray) in ICB treated mice. E. CREB5 protein expression in B16 cells expressing control or *CREB5* ORF. F. Tumor growth over time in untreated mice and G. Survival curve in ICB treated mice for *CREB5* ORF tumors (red) relative to control (Gray). H. B16 Tumor growth over time in NSG (Black), WT (Grey), WT + ICB (Red), and WT + ICB + aCD8 (Purple) mice. I. *In vivo* competition assay showing log2 fold change for *Creb5* asgRNA1 relative to control in B16 tumors. J. *Creb5* transcript abundance in B16 and YUMMER cells K. *Creb5* transcript abundance in YUMMER control *Creb5* KO cells (Blue) relative to control (Gray) and protein expression. L. Results of *in vivo* competition assay showing log2 fold change for *Creb5* KO sgRNA relative to control in YUMMER tumors. M. *Creb5* transcript abundance in YUMMER *Creb5* asgRNA cells (Red) relative to control (Gray). N. Results of *in vivo* competition assay showing log2 fold change for *Creb5* asgRNA relative to control in YUMMER tumors.

**Fig. S3.**
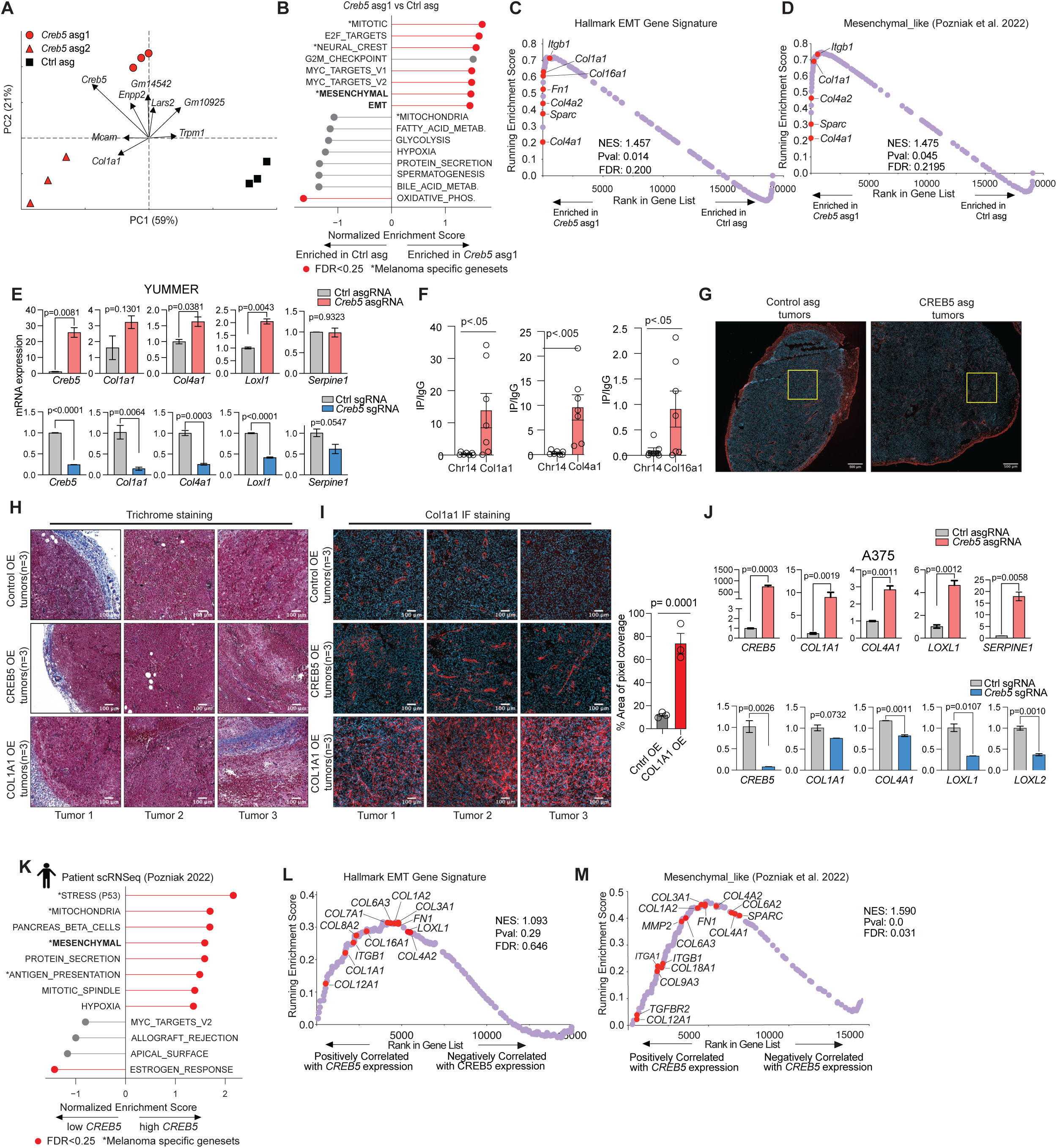
CREB5 promotes expression of collagen and collagen-associated genes in cancer cells. A. PCA of transcriptional profiling of *Creb5* asg1, *Creb5* asg2, and control B16 cells. The top genes driving variation between the samples are plotted as vectors. B. Bar graph showing the gene set pathways enriched in *Creb5* asg1 cells relative to control B16 cells. C-D. Mountain plot showing enrichment score for the C. EMT gene set and D. Mesenchymal-like gene set in *Creb5* asg1 cells relative to control B16 cells. Gene symbols and ranks for collagen and collagen-related genes are marked in red. E. Transcript abundance for collagen and collagen-stabilizing factors in *Creb5* asgRNA cells (Red, top) or *Creb5* sgRNA cells (Blue, bottom) relative to control (Grey) in YUMMER cells. F. Transcript abundance of FOXA1 motifs in the promoter region of *Col1a1*, *Col4a1*, and *Col16a1*. G. Zoom out IF image (500um) of control OE tumors (Left) and Creb5 OE tumors (Right) shown in Fig 3G. H. Representative images of trichrome staining in control OE (Top), *Creb5* OE (Middle) and *Col1a1* OE (Bottom) tumors. I. Col1a1 IF staining of control OE (Top), *Creb5* OE (Middle) and *Col1a1* OE (Bottom) tumors and quantification on left. J. Transcript abundance for collagen and collagen-stabilizing factors in *Creb5* asgRNA cells (Red, top) or *Creb5* sgRNA cells (Blue, bottom) relative to control (Grey) in A375 cells. K. Bar graph showing the gene set pathways correlated with *CREB5* expression in human melanoma patient samples. L-M. Mountain plot showing enrichment score for the H. EMT gene set and I. Mesenchymal-like gene set for correlation with *CREB5* expression in human melanoma patient samples. Gene symbols and ranks for collagen and collagen-related genes are marked in red.

**Fig. S4.**
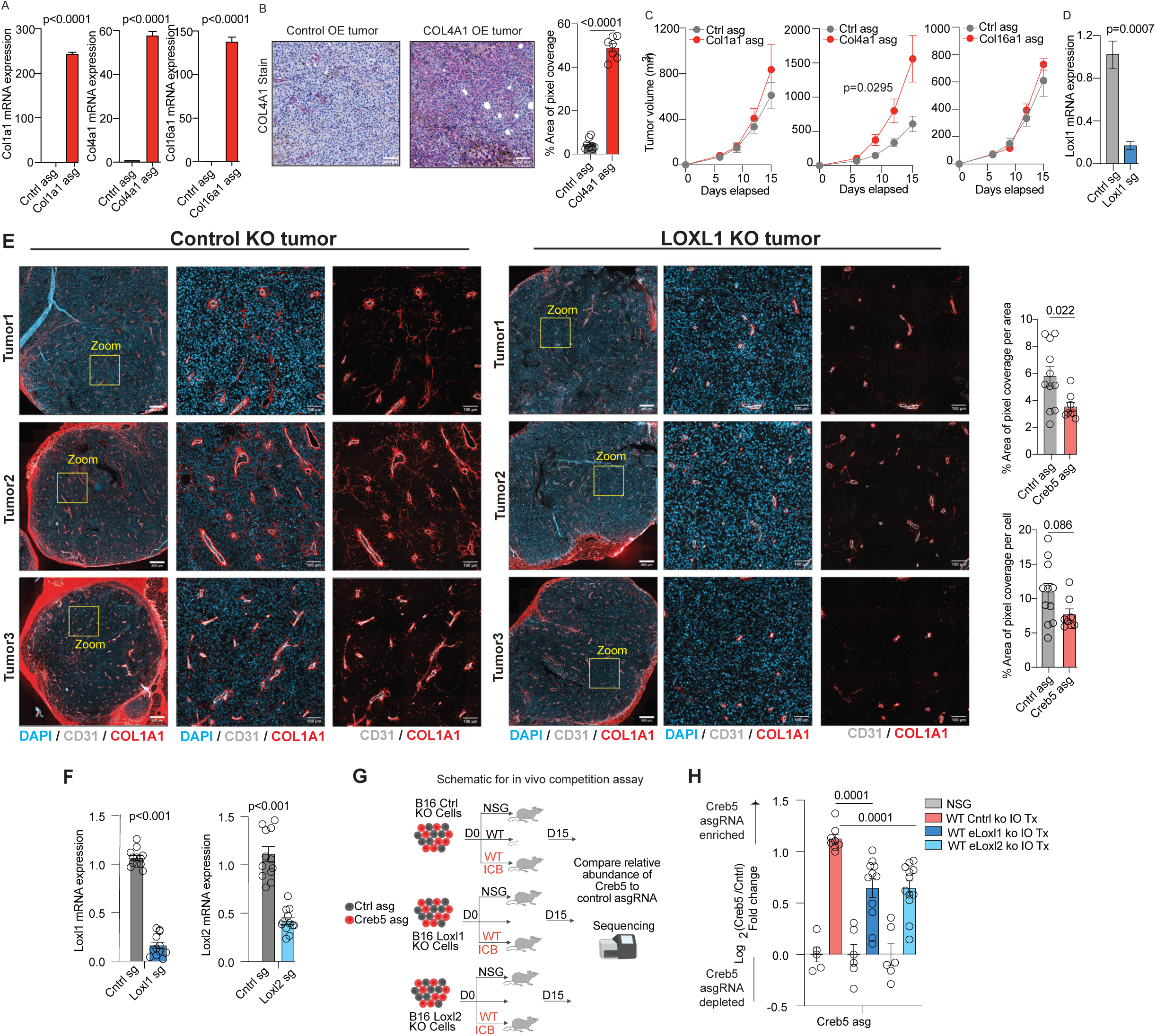
Tumor-intrinsic collagen matrix deposition promotes ICB resistance. A. Transcript abundance of *Col1a1* (Left), *Col4a1* (Center), *Col16a1* (Right) in B16 cells with respective target asgRNA (Red) relative to control (Grey). B. COL4A1 IHC staining on control OE (Left) and Creb5 OE tumors (Center), and quantification of pixel coverage over area (Right). C. B16 Tumor growth over time of *Col1a1* asg (Left), *Col4a1* asg (Center) and *Col16a1* asg (Right) in red relative to control asg in gray in WT mice. D. Transcript abundance of *Loxl1* in B16 *Loxl1* KO sgRNA cells (Blue) relative to control (Gray). E. IF staining of COL1A1 (Red), CD31 (White), and DAPI (Blue) in control ko tumors (Left) and Loxl1 ko tumors (Center) and quantification of pixel coverage of COL1A1 per area (Right, top) and per cell (Right, bottom). F. Transcript abundance of *Loxl1* in B16 *Loxl1* KO sgRNA cells (Blue) relative to control (Gray) on left and, transcript abundance of *Loxl2* in B16 *Loxl2* KO sgRNA cells (Blue) relative to control (Grey) on Right. G. Schematic of *in vivo* competition assay. H. Results of *in vivo* competition assay showing log2 fold change for *Creb5* asgRNA relative to control in B16 control KO tumors (Left) or B16 *Loxl1* KO tumors (Center) and B16 *Loxl2* ko (Right).

**Fig. S5.**
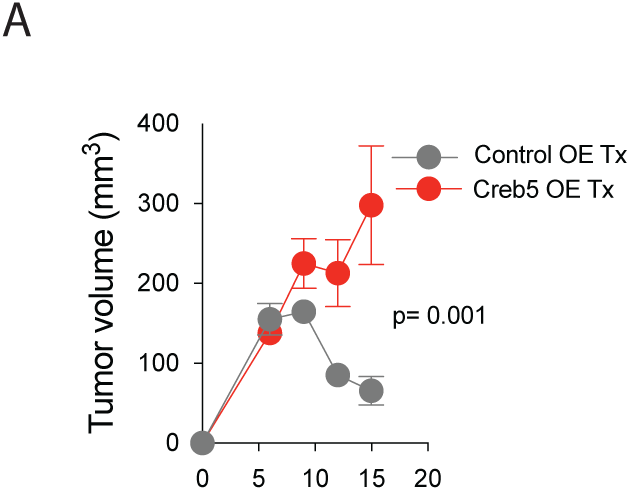
CREB5 promotes immunosuppressive tumor microenvironment. A. Tumor growth over time in ICB treated mice for Creb5 OE tumors (Red) relative to control (Gray).

**Fig. S6.**
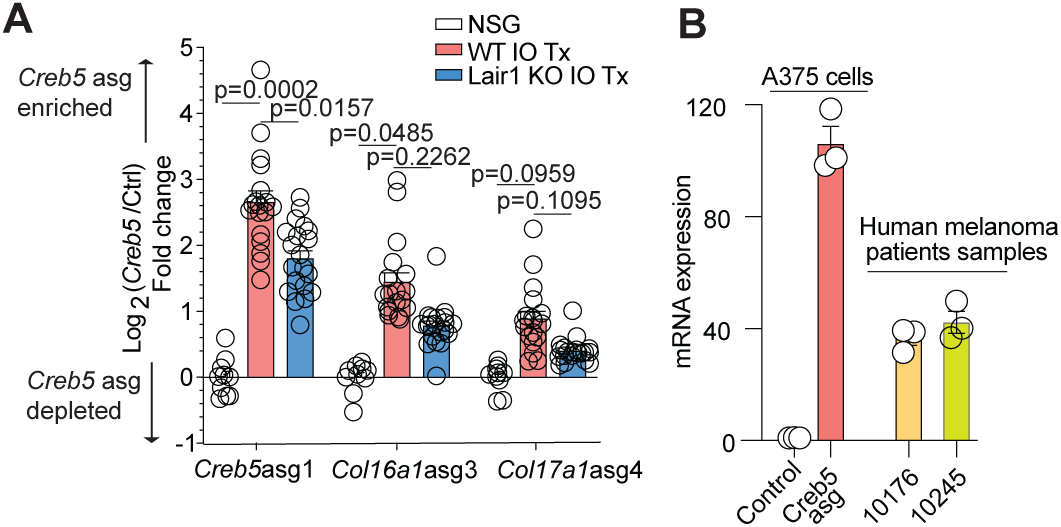
Collagen sensing by LAIR1 is required for CREB5-mediated immunotherapy resistance. A. Results of *in vivo* competition assay showing log2 fold change for *Creb5* asgRNA1 (Left), *Col16a1* asgRNA3 (Middle), and *Col17a1* asgRNA4 (Right) relative to control in B16 tumors. B. Transcript abundance of *CREB5* in A375 and patient samples.

**Table S6:**
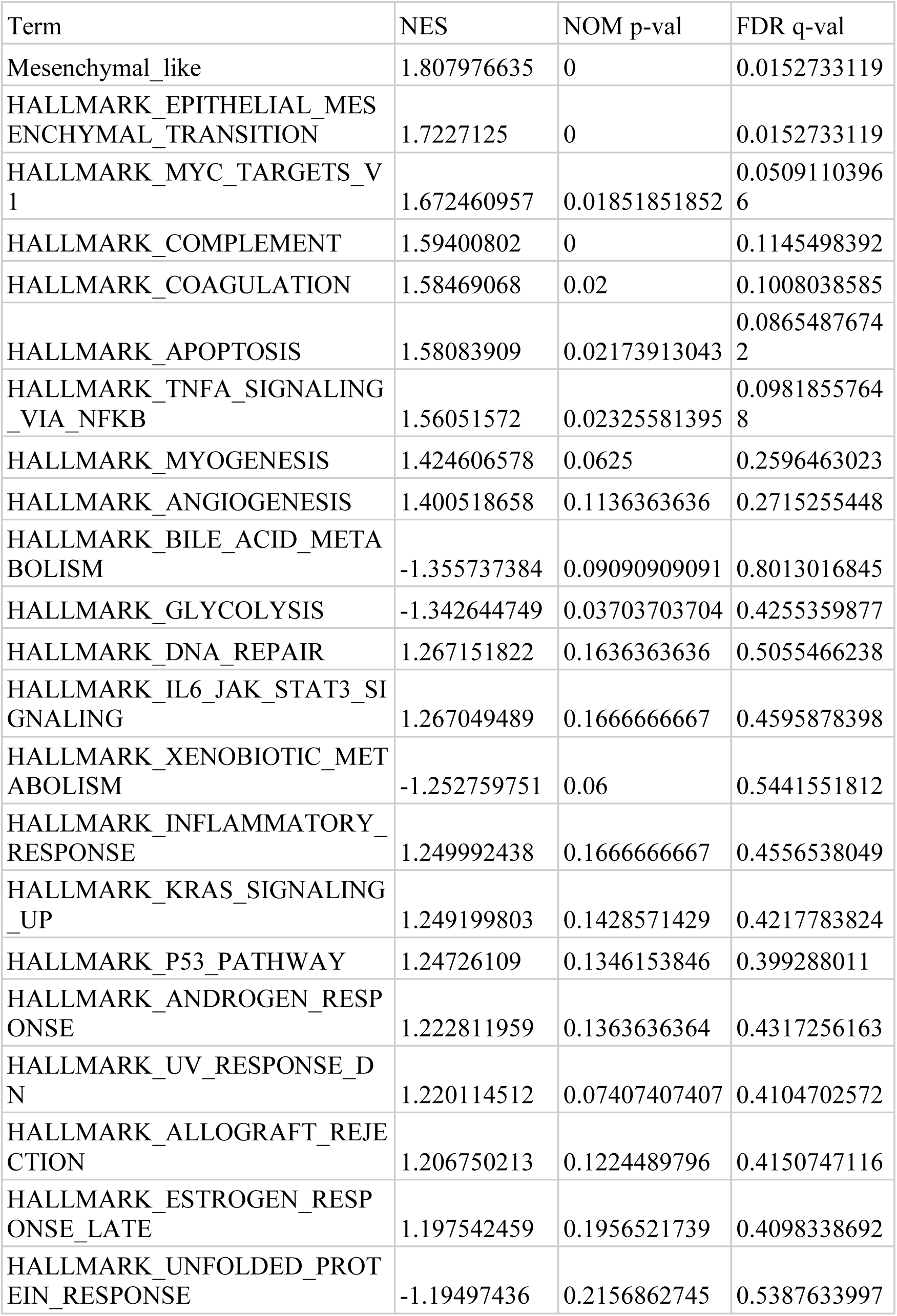
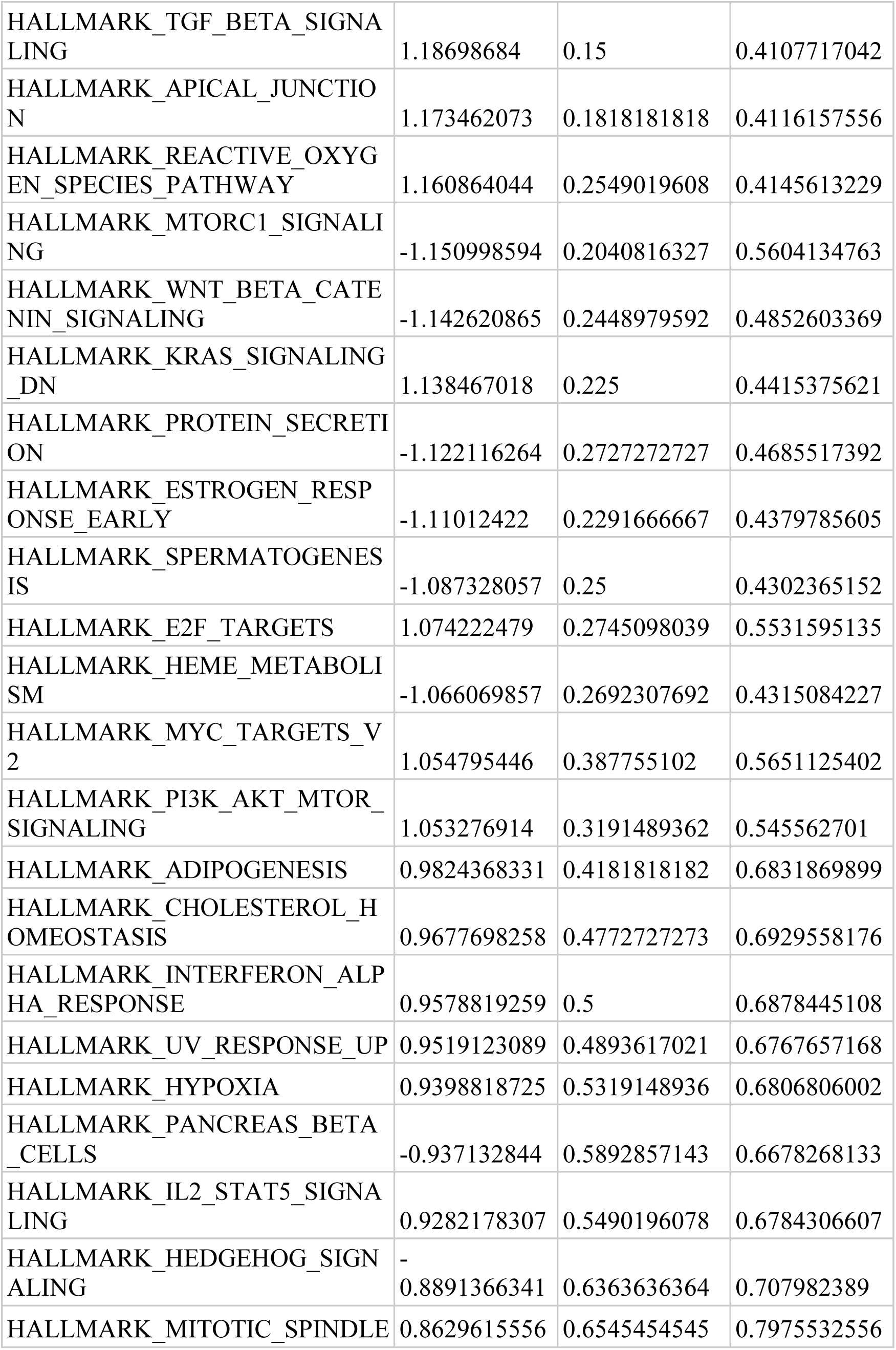
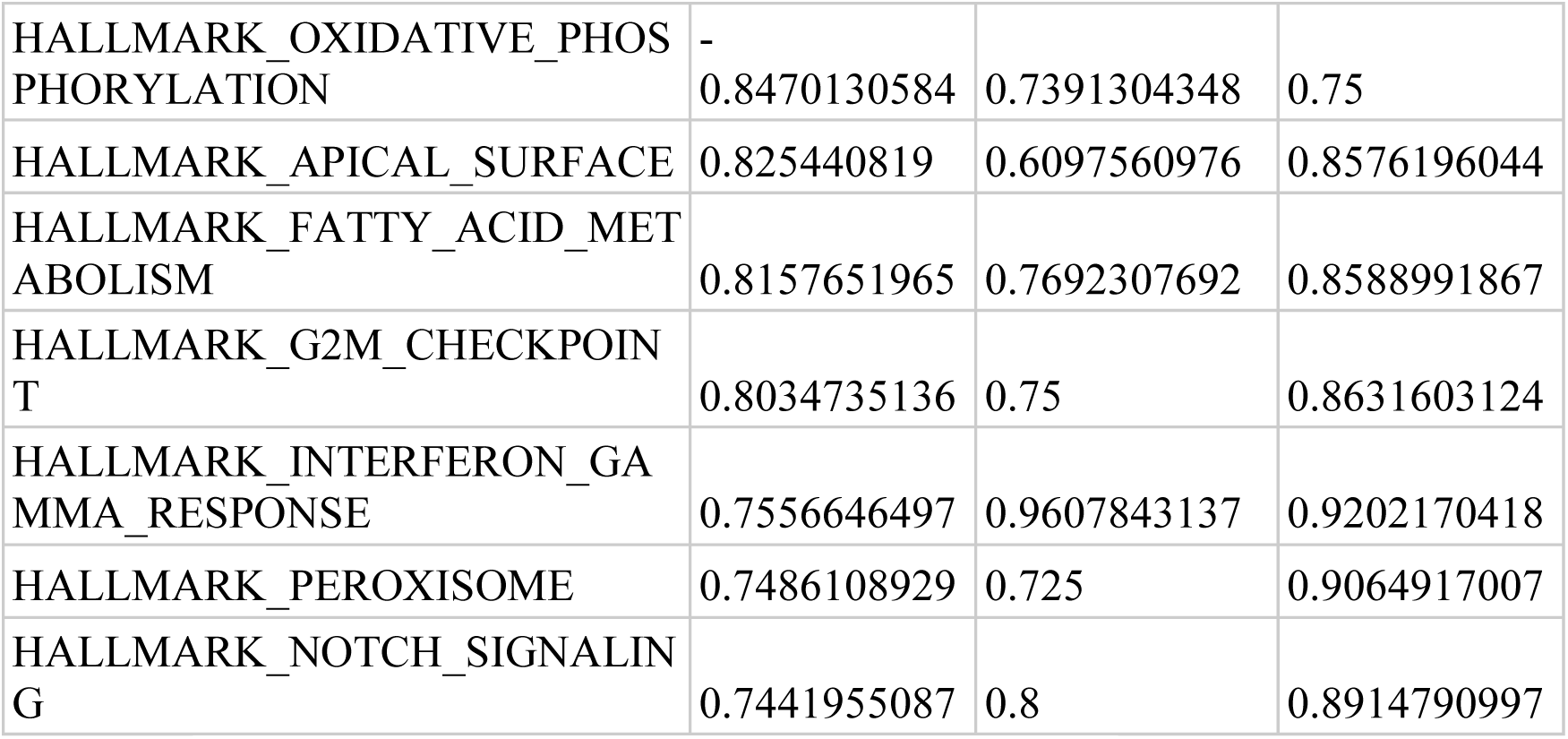
Bulk RNA-seq CREB5 sg2 vs CTRL GSEA results.

**Table S8:**
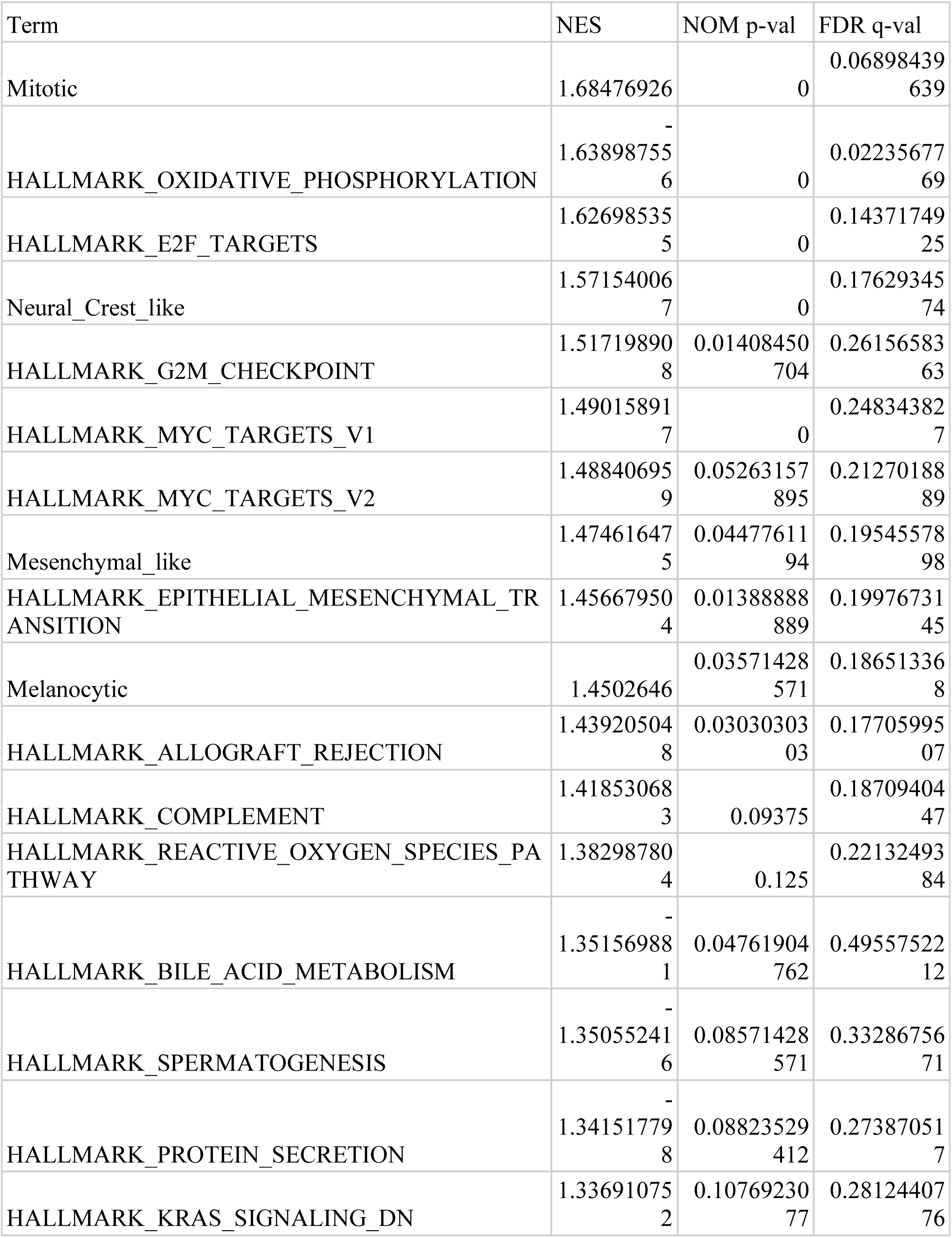
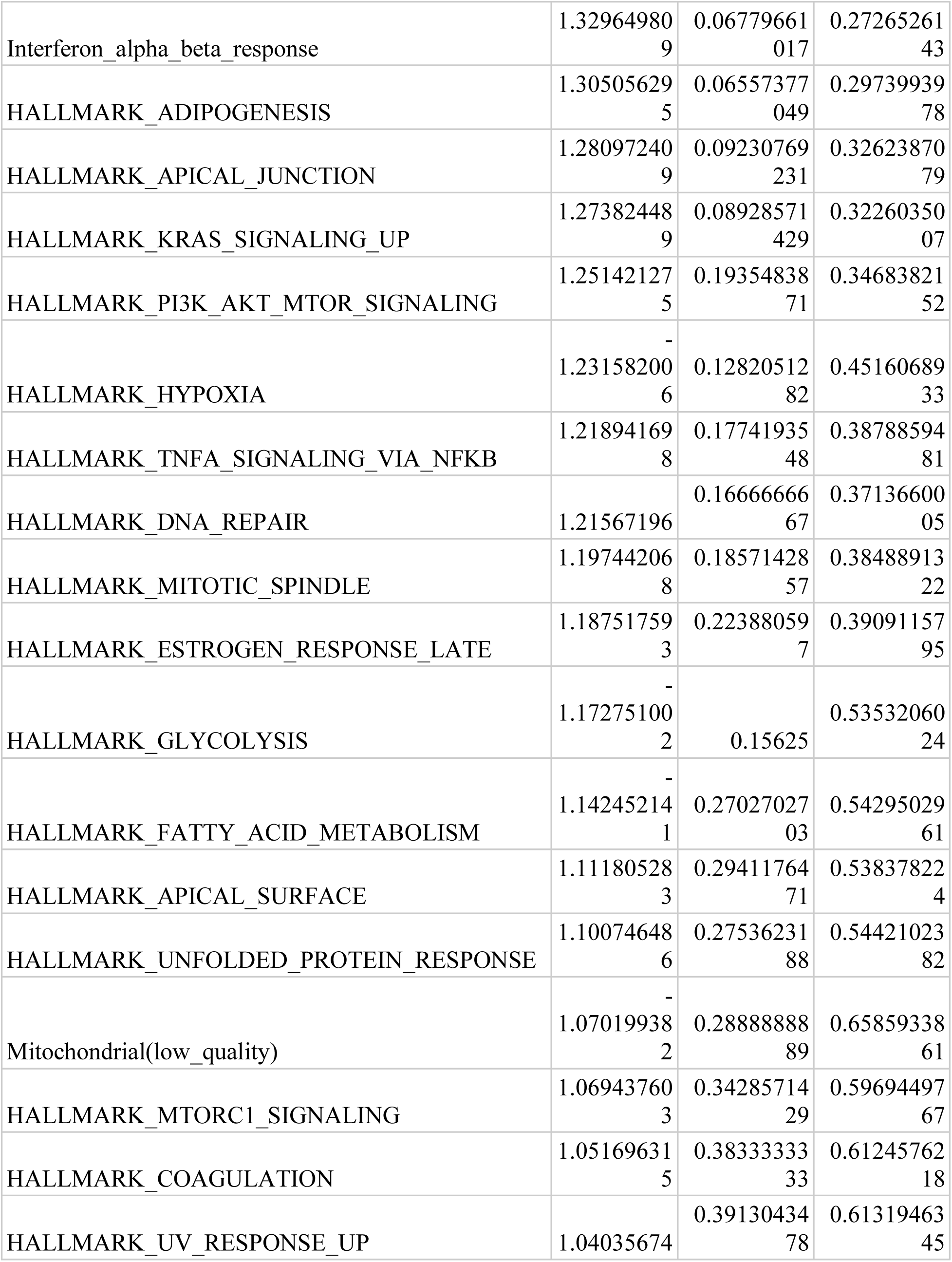
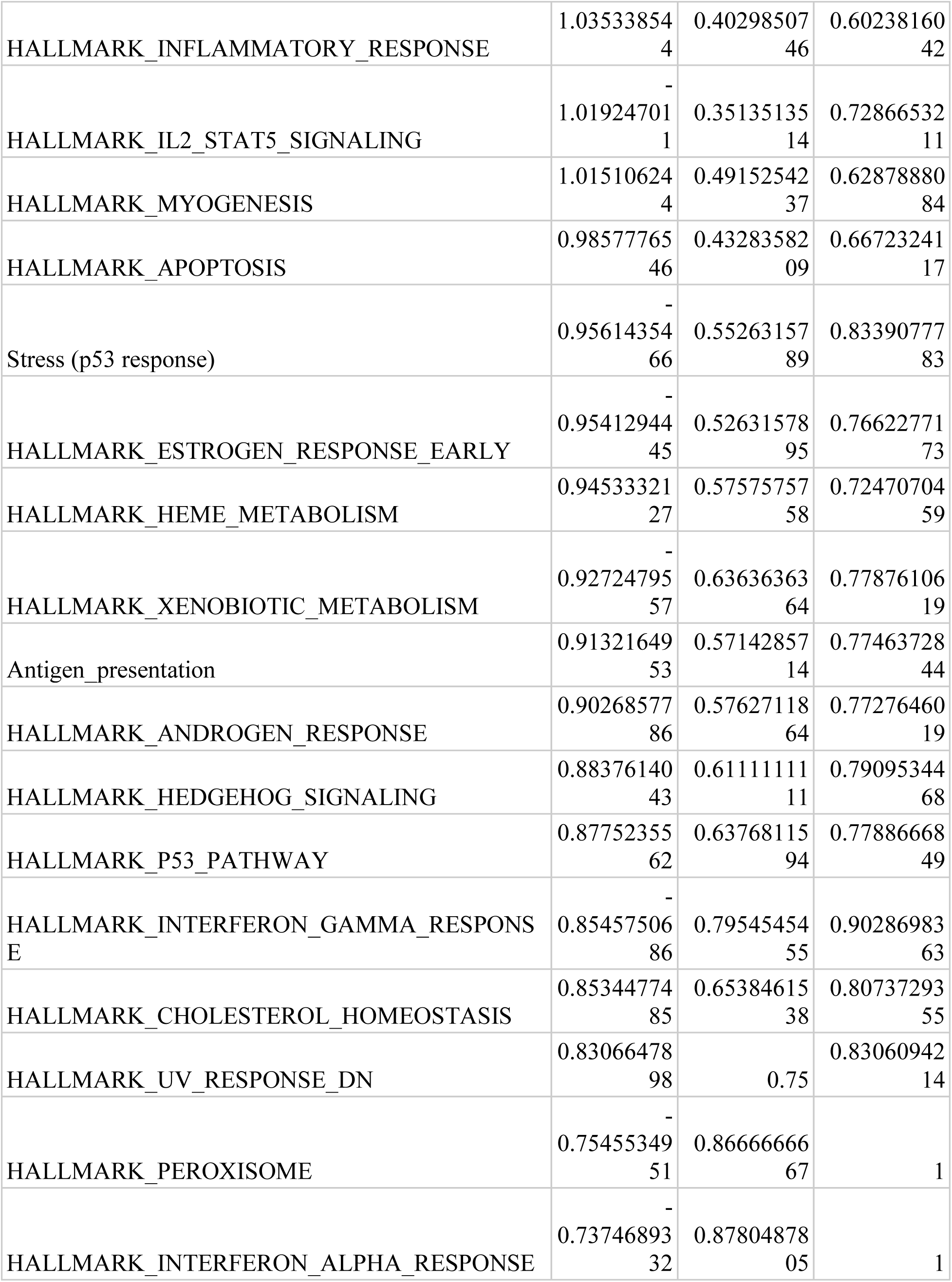
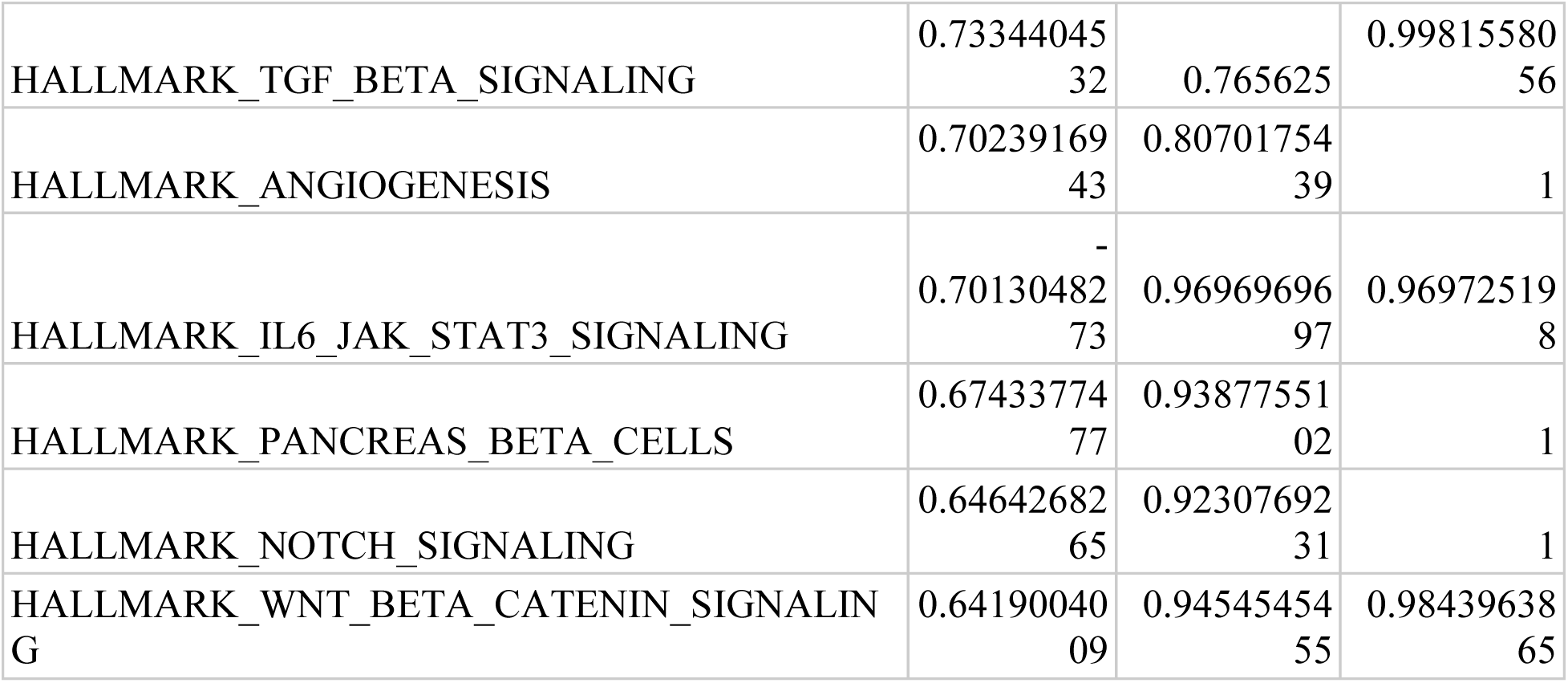
Bulk RNA-seq *CREB5* sg1 vs CTRL GSEA results.

**Table S9:** Jerby-Anon scRNA-seq - CREB5 correlation GSEA.

| Term | NES | NOM p-val | FDR q-val |
| --- | --- | --- | --- |
| Mesenchymal_like | 2.54326474<br>8 | 0 | 0 |
| Stress (p53 response) | 1.71370981<br>7 | 0 | 0.02105263<br>158 |
| HALLMARK_COAGULATION | 1.60332325<br>4 | 0 | 0.01403508<br>772 |
| Mitochondrial(low_quality) | 1.27186129<br>5 | 0.11111111<br>11 | 0.19473684<br>21 |
| HALLMARK_EPITHELIAL_MESENCHYMAL_TRANSITION | 1.23434789<br>3 | 0 | 0.21052631<br>58 |
| HALLMARK_KRAS_SIGNALING_UP | 1.17517794<br>1 | 0.16666666<br>67 | 0.28070175<br>44 |
| HALLMARK_MITOTIC_SPINDLE | 1.16280852<br>7 | 0 | 0.25563909<br>77 |
| HALLMARK_PROTEIN_SECRETION | 1.07533321<br>2 | 0.23076923<br>08 | 0.43157894<br>74 |
| HALLMARK_UV_RESPONSE_DN | 1.06010195 | 0.33333333<br>33 | 0.42105263<br>16 |
| Neural_Crest_like | 1 | 1 | 0.54947368<br>42 |
| HALLMARK_ANGIOGENESIS | 0.76332534<br>54 | 0.90476190<br>48 | 1 |
| HALLMARK_WNT_BETA_CATENIN_SIGNALING | 0.57013834<br>89 | 1 | 1 |
| HALLMARK_BILE_ACID_METABOLISM | -<br>0.69625381<br>42 | 0.94382022<br>47 | 0.96454033<br>77 |
| HALLMARK_HEME_METABOLISM | -<br>0.70573370<br>35 | 1 | 0.98014112<br>08 |
| HALLMARK_SPERMATOGENESIS | -<br>0.71165157<br>07 | 0.96511627<br>91 | 0.99702313<br>95 |
| Antigen_presentation | -<br>0.72291520<br>29 | 0.98958333<br>33 | 1 |
| HALLMARK_TGF_BETA_SIGNALING | -<br>0.72663608<br>96 | 0.85542168<br>67 | 1 |
| HALLMARK_PI3K_AKT_MTOR_SIGNALING | -<br>0.73058895<br>11 | 0.93103448<br>28 | 1 |
| HALLMARK_FATTY_ACID_METABOLISM | -<br>0.74130617<br>8 | 0.92473118<br>28 | 1 |
| HALLMARK_ANDROGEN_RESPONSE | -<br>0.77847187<br>27 | 0.87058823<br>53 | 1 |
| HALLMARK_COMPLEMENT | -<br>0.77879652<br>14 | 0.88297872<br>34 | 1 |
| HALLMARK_MYOGENESIS | -<br>0.80208327<br>49 | 0.89247311<br>83 | 1 |
| HALLMARK_CHOLESTEROL_HOMEOSTASIS | -<br>0.80646969<br>79 | 0.83529411<br>76 | 1 |
| HALLMARK_NOTCH_SIGNALING | -<br>0.82106227<br>27 | 0.7375 | 1 |
| HALLMARK_GLYCOLYSIS | -<br>0.83710328<br>32 | 0.84444444<br>44 | 1 |
| Mitotic | -<br>0.84075910<br>48 | 0.90816326<br>53 | 1 |
| HALLMARK_INTERFERON_ALPHA_RESPONSE | -<br>0.84565871<br>7 | 0.72941176<br>47 | 1 |
| HALLMARK_HEDGEHOG_SIGNALING | -<br>0.86239978<br>7 | 0.67142857<br>14 | 1 |
| HALLMARK_E2F_TARGETS | -<br>0.87147898<br>63 | 0.77659574<br>47 | 1 |
| HALLMARK_APOPTOSIS | -<br>0.87871974<br>6 | 0.72527472<br>53 | 1 |
| HALLMARK_PANCREAS_BETA_CELLS | -<br>0.88039403<br>72 | 0.72619047<br>62 | 1 |
| HALLMARK_ESTROGEN_RESPONSE_EARLY | -<br>0.88337508<br>9 | 0.75510204<br>08 | 1 |
| HALLMARK_G2M_CHECKPOINT | -<br>0.88864132<br>72 | 0.72164948<br>45 | 1 |
| HALLMARK_APICAL_JUNCTION | -<br>0.90662332<br>84 | 0.70526315<br>79 | 1 |
| HALLMARK_HYPOXIA | -<br>0.91093086<br>02 | 0.69791666<br>67 | 1 |
| HALLMARK_MYC_TARGETS_V1 | -<br>0.95083136<br>41 | 0.65217391<br>3 | 1 |
| Interferon_alpha_beta_response | -<br>0.96546697<br>56 | 0.57575757<br>58 | 1 |
| HALLMARK_KRAS_SIGNALING_DN | -<br>0.97850798<br>82 | 0.5 | 1 |
| HALLMARK_ESTROGEN_RESPONSE_LATE | -<br>0.99695112<br>73 | 0.5 | 1 |
| HALLMARK_DNA_REPAIR | -<br>1.00990619<br>6 | 0.48913043<br>48 | 1 |
| HALLMARK_IL2_STAT5_SIGNALING | -<br>1.04123087<br>9 | 0.37234042<br>55 | 0.98065567<br>3 |
| HALLMARK_PEROXISOME | -<br>1.04164937<br>2 | 0.35632183<br>91 | 1 |
| HALLMARK_INTERFERON_GAMMA_RESPONSE | -<br>1.06727282<br>8 | 0.36956521<br>74 | 0.95649486<br>81 |
| HALLMARK_APICAL_SURFACE | -<br>1.07054572<br>5 | 0.32530120<br>48 | 0.99312851<br>78 |
| Stress (hypoxia response) | -<br>1.10734883<br>3 | 0.27956989<br>25 | 0.87415884<br>93 |
| HALLMARK_UV_RESPONSE_UP | -<br>1.12616619<br>6 | 0.25555555<br>56 | 0.83582149<br>56 |
| HALLMARK_MTORC1_SIGNALING | -<br>1.12838673<br>2 | 0.23157894<br>74 | 0.88790590<br>27 |
| HALLMARK_P53_PATHWAY | -<br>1.15636493<br>6 | 0.14285714<br>29 | 0.77157598<br>5 |
| HALLMARK_TNFA_SIGNALING_VIA_NFKB | -<br>1.17138196<br>9 | 0.16494845<br>36 | 0.74953095<br>68 |
| HALLMARK_UNFOLDED_PROTEIN_RESPONSE | -<br>1.19184351<br>5 | 0.16279069<br>77 | 0.69221388<br>37 |
| HALLMARK_MYC_TARGETS_V2 | -<br>1.21676514 | 0.20987654<br>32 | 0.61432145<br>09 |
| HALLMARK_IL6_JAK_STAT3_SIGNALING | -<br>1.22965216<br>5 | 0.15957446<br>81 | 0.62828330<br>21 |
| HALLMARK_XENOBIOTIC_METABOLISM | -<br>1.23557410<br>8 | 0.07777777<br>778 | 0.68528544<br>63 |
| HALLMARK_ALLOGRAFT_REJECTION | -<br>1.25540615<br>2 | 0.05154639<br>175 | 0.66135084<br>43 |
| HALLMARK_ADIPOGENESIS | -<br>1.26839674<br>4 | 0.05208333<br>333 | 0.70367729<br>83 |
| HALLMARK_INFLAMMATORY_RESPONSE | -<br>1.29864711<br>6 | 0.05434782<br>609 | 0.65253283<br>3 |
| HALLMARK_REACTIVE_OXYGEN_SPECIES_PATHWAY | -<br>1.33524804<br>7 | 0.05263157<br>895 | 0.58786741<br>71 |
| HALLMARK_OXIDATIVE_PHOSPHORYLATION | -<br>1.42213452<br>4 | 0.02247191<br>011 | 0.31744840<br>53 |
| Melanocytic | -<br>1.88483765<br>6 | 0 | 0 |

**Table S10:** Pozniak scRNA-seq - CREB5 correlation GSEA.

| Term | NES | NOM p-val | FDR q-val |
| --- | --- | --- | --- |
| Stress (p53 response) | 2.1627697<br>99 | 0 | 0 |
| Mitochondrial(low_quality) | 1.6961126<br>65 | 0 | 0.009578191<br>195 |
| HALLMARK_PANCREAS_BETA_CELLS | 1.6919987<br>12 | 0 | 0.009578191<br>195 |
| Mesenchymal_like | 1.5901684<br>93 | 0 | 0.031129121<br>39 |
| HALLMARK_PROTEIN_SECRETION | 1.5885421<br>04 | 0 | 0.024903297<br>11 |
| Antigen_presentation | 1.4887182<br>13 | 0 | 0.060661877<br>57 |
| HALLMARK_ESTROGEN_RESPONSE_LATE | -<br>1.4256158<br>09 | 0 | 0.163742690<br>1 |
| HALLMARK_MITOTIC_SPINDLE | 1.3888336<br>68 | 0.01010101<br>01 | 0.135462989<br>8 |
| HALLMARK_HYPOXIA | 1.3620511<br>04 | 0.01 | 0.168815619<br>8 |
| HALLMARK_PI3K_AKT_MTOR_SIGNALING | 1.2960787<br>22 | 0.06060606<br>061 | 0.329915474<br>5 |
| HALLMARK_WNT_BETA_CATENIN_SIGNALING | 1.2900344<br>98 | 0.12087912<br>09 | 0.315122490<br>3 |
| Melanocytic | 1.2889976<br>33 | 0.02040816<br>327 | 0.290828714<br>5 |
| HALLMARK_MYC_TARGETS_V1 | 1.2779972<br>65 | 0.03061224<br>49 | 0.299318474<br>9 |
| HALLMARK_ANGIOGENESIS | 1.2634958<br>76 | 0.17977528<br>09 | 0.316817093<br>4 |
| HALLMARK_GLYCOLYSIS | 1.2404525<br>54 | 0.09 | 0.359182169<br>8 |
| HALLMARK_TGF_BETA_SIGNALING | 1.2278213<br>58 | 0.16129032<br>26 | 0.372910910<br>5 |
| HALLMARK_APOPTOSIS | 1.2225407<br>92 | 0.10101010<br>1 | 0.368760361 |
| HALLMARK_ESTROGEN_RESPONSE_EARLY | 1.2198989<br>09 | 0.09 | 0.354956497<br>2 |
| HALLMARK_ANDROGEN_RESPONSE | 1.2087517<br>34 | 0.14141414<br>14 | 0.376742187 |
| HALLMARK_MYOGENESIS | 1.189520068 | 0.11 | 0.4204321819 |
| HALLMARK_APICAL_SURFACE | -1.166111299 | 0.2307692308 | 0.350877193 |
| HALLMARK_IL2_STAT5_SIGNALING | 1.16115259 | 0.112244898 | 0.50046049 |
| HALLMARK_INTERFERON_ALPHA_RESPONSE | 1.132834086 | 0.3195876289 | 0.5815330369 |
| HALLMARK_APICAL_JUNCTION | 1.118666122 | 0.2323232323 | 0.6086505132 |
| HALLMARK_UV_RESPONSE_DN | 1.099431608 | 0.2929292929 | 0.6525663306 |
| HALLMARK_EPITHELIAL_MESENCHYMAL_TRANSITION | 1.093406603 | 0.29 | 0.6457297231 |
| HALLMARK_KRAS_SIGNALING_UP | 1.091369317 | 0.28 | 0.629861853 |
| HALLMARK_TNFA_SIGNALING_VIA_NFKB | 1.070016256 | 0.33 | 0.6977343894 |
| HALLMARK_INTERFERON_GAMMA_RESPONSE | 1.05496945 | 0.3636363636 | 0.7325542525 |
| HALLMARK_SPERMATOGENESIS | 1.025197168 | 0.4736842105 | 0.8196195037 |
| Interferon_alpha_beta_response | 1.024628579 | 0.42 | 0.7936687394 |
| HALLMARK_HEDGEHOG_SIGNALING | 1.022884256 | 0.3720930233 | 0.7745563947 |
| HALLMARK_ALLOGRAFT_REJECTION | -1 | 1 | 0.6549707602 |
| HALLMARK_P53_PATHWAY | 0.9618297544 | 0.6060606061 | 0.9667793629 |
| HALLMARK_INFLAMMATORY_RESPONSE | 0.939379001 | 0.6326530612 | 1 |
| HALLMARK_MTORC1_SIGNALING | 0.9236186307 | 0.6464646465 | 1 |
| HALLMARK_CHOLESTEROL_HOMEOSTASIS | 0.921417511 | 0.6526315789 | 1 |
| HALLMARK_BILE_ACID_METABOLISM | 0.9162364231 | 0.6914893617 | 0.9996894982 |
| HALLMARK_COAGULATION | 0.9015154246 | 0.7395833333 | 1 |
| HALLMARK_PEROXISOME | 0.8917090<br>026 | 0.71739130<br>43 | 1 |
| HALLMARK_FATTY_ACID_METABOLISM | 0.8803937<br>452 | 0.69696969<br>7 | 1 |
| HALLMARK_DNA_REPAIR | 0.8753284<br>766 | 0.73 | 0.994167127<br>2 |
| HALLMARK_HEME_METABOLISM | 0.8493522<br>836 | 0.78 | 1 |
| HALLMARK_IL6_JAK_STAT3_SIGNALING | 0.8241805<br>369 | 0.75531914<br>89 | 1 |
| HALLMARK_OXIDATIVE_PHOSPHORYLATION | 0.8171025<br>869 | 0.84 | 1 |
| HALLMARK_UNFOLDED_PROTEIN_RESPONSE | 0.8146290<br>124 | 0.82653061<br>22 | 1 |
| HALLMARK_G2M_CHECKPOINT | 0.8127765<br>494 | 0.87878787<br>88 | 0.989383613<br>3 |
| HALLMARK_MYC_TARGETS_V2 | -<br>0.8053661<br>337 | 1 | 0.865497076 |
| HALLMARK_ADIPOGENESIS | 0.8017687<br>598 | 0.91 | 0.985915147 |
| HALLMARK_NOTCH_SIGNALING | 0.7971386<br>576 | 0.77647058<br>82 | 0.971561741<br>7 |
| HALLMARK_KRAS_SIGNALING_DN | 0.7806182<br>184 | 0.87628865<br>98 | 0.975956545<br>4 |
| HALLMARK_XENOBIOTIC_METABOLISM | 0.7752630<br>904 | 0.86868686<br>87 | 0.962408669<br>5 |
| HALLMARK_COMPLEMENT | 0.7637703<br>519 | 0.92 | 0.954887020<br>2 |
| HALLMARK_UV_RESPONSE_UP | 0.7568383<br>423 | 0.89898989<br>9 | 0.942110886 |
| HALLMARK_REACTIVE_OXYGEN_SPECIES_PATHWAY | 0.7043427<br>684 | 0.89247311<br>83 | 0.963641157<br>3 |
| HALLMARK_E2F_TARGETS | 0.5177871<br>377 | 1 | 0.998158040<br>2 |

**Table S12:** Liu RNA-seq - CREB5 correlation GSEA.

| Term | NES | NOM<br>p-val | FDR q-val |
| --- | --- | --- | --- |
| HALLMARK_G2M_CHECKPOINT | 1.86672<br>3652 | 0 | 0 |
| Mitochondrial(low_quality) | 1.83544<br>4684 | 0 | 0 |
| Neural_Crest_like | 1.82349<br>1944 | 0 | 0 |
| HALLMARK_TGF_BETA_SIGNALING | 1.75437<br>1762 | 0 | 0 |
| Mitotic | 1.71122<br>0345 | 0 | 0.001921097<br>77 |
| Mesenchymal_like | 1.70788<br>3409 | 0 | 0.001600914<br>808 |
| HALLMARK_PROTEIN_SECRETION | 1.66060<br>6826 | 0 | 0.001372212<br>693 |
| HALLMARK_E2F_TARGETS | 1.64406<br>6816 | 0 | 0.001200686<br>106 |
| HALLMARK_MITOTIC_SPINDLE | 1.62401<br>9983 | 0 | 0.002134553<br>078 |
| Stress (p53 response) | 1.58651<br>0164 | 0.01030<br>927835 | 0.002881646<br>655 |
| HALLMARK_MYC_TARGETS_V1 | 1.54793<br>4789 | 0 | 0.006112583<br>814 |
| HALLMARK_UV_RESPONSE_DN | 1.51174<br>8521 | 0 | 0.008805031<br>447 |
| HALLMARK_WNT_BETA_CATENIN_SIGNALING | 1.51165<br>1371 | 0.02105<br>263158 | 0.008127721<br>335 |
| HALLMARK_ANDROGEN_RESPONSE | 1.41301<br>7637 | 0 | 0.026072041<br>17 |
| HALLMARK_HEDGEHOG_SIGNALING | 1.40272<br>4628 | 0.03296<br>703297 | 0.030097198<br>4 |
| HALLMARK_EPITHELIAL_MESENCHYMAL_TRANSITION | 1.36203<br>5738 | 0.01 | 0.053430531<br>73 |
| HALLMARK_MTORC1_SIGNALING | 1.29495<br>866 | 0.02 | 0.124871355<br>1 |
| HALLMARK_PI3K_AKT_MTOR_SIGNALING | 1.27484<br>1857 | 0.09 | 0.152086906<br>8 |
| HALLMARK_COMPLEMENT | 1.23991<br>0309 | 0.06 | 0.207276338<br>4 |
| HALLMARK_UNFOLDED_PROTEIN_RESPONSE | 1.224423506 | 0.13 | 0.2305317324 |
| HALLMARK_DNA_REPAIR | 1.211419931 | 0.1 | 0.2511149228 |
| HALLMARK_PANCREAS_BETA_CELLS | 1.202680108 | 0.1632653061 | 0.2637143303 |
| HALLMARK_ANGIOGENESIS | 1.198641538 | 0.2444444444 | 0.2614363487 |
| HALLMARK_NOTCH_SIGNALING | 1.197203795 | 0.2065217391 | 0.2521440823 |
| HALLMARK_KRAS_SIGNALING_UP | 1.192966017 | 0.06 | 0.2520480274 |
| HALLMARK_REACTIVE_OXYGEN_SPECIES_PATHWAY | 1.180845413 | 0.1894736842 | 0.2781897348 |
| HALLMARK_APICAL_JUNCTION | 1.174575919 | 0.11 | 0.2910107363 |
| HALLMARK_SPERMATOGENESIS | 1.162228231 | 0.2121212121 | 0.314922813 |
| Stress (hypoxia response) | 1.13223478 | 0.22 | 0.3954811616 |
| HALLMARK_APOPTOSIS | 1.116373682 | 0.22 | 0.4341680961 |
| HALLMARK_HEME_METABOLISM | 1.085641869 | 0.31 | 0.5410059204 |
| HALLMARK_IL2_STAT5_SIGNALING | 1.081425996 | 0.27 | 0.5406089194 |
| HALLMARK_CHOLESTEROL_HOMEOSTASIS | 1.017378546 | 0.4583333333 | 0.7949269713 |
| HALLMARK_TNFA_SIGNALING_VIA_NFKB | 0.9874588213 | 0.53 | 0.8896377762 |
| HALLMARK_INFLAMMATORY_RESPONSE | 0.9419806363 | 0.67 | 1 |
| HALLMARK_COAGULATION | 0.9220428431 | 0.73 | 1 |
| HALLMARK_FATTY_ACID_METABOLISM | 0.8925917808 | 0.72 | 1 |
| HALLMARK_MYC_TARGETS_V2 | 0.8685065092 | 0.7395833333 | 1 |
| HALLMARK_P53_PATHWAY | 0.8484897069 | 0.85 | 1 |
| Antigen_presentation | 0.84782<br>3243 | 0.9 | 1 |
| HALLMARK_GLYCOLYSIS | 0.81325<br>89396 | 0.9 | 1 |
| HALLMARK_IL6_JAK_STAT3_SIGNALING | 0.80367<br>4116 | 0.86 | 1 |
| HALLMARK_ESTROGEN_RESPONSE_EARLY | 0.78974<br>80059 | 0.91 | 1 |
| HALLMARK_HYPOXIA | 0.77290<br>70977 | 0.96 | 1 |
| HALLMARK_OXIDATIVE_PHOSPHORYLATION | 0.77208<br>66295 | 0.95 | 1 |
| HALLMARK_XENOBIOTIC_METABOLISM | 0.72593<br>51709 | 0.98 | 1 |
| HALLMARK_ALLOGRAFT_REJECTION | 0.72314<br>86272 | 0.99 | 1 |
| HALLMARK_MYOGENESIS | 0.70985<br>17103 | 0.99 | 1 |
| HALLMARK_ADIPOGENESIS | 0.65650<br>4269 | 0.98 | 1 |
| HALLMARK_UV_RESPONSE_UP | 0.65114<br>14755 | 1 | 1 |
| HALLMARK_PEROXISOME | 0.63570<br>51403 | 0.96969<br>69697 | 1 |
| HALLMARK_INTERFERON_GAMMA_RESPONSE | 0.60801<br>96064 | 0.98 | 1 |
| HALLMARK_APICAL_SURFACE | 0.60122<br>99551 | 0.94623<br>65591 | 1 |
| HALLMARK_KRAS_SIGNALING_DN | 0.59453<br>90242 | 1 | 1 |
| HALLMARK_BILE_ACID_METABOLISM | 0.58532<br>61378 | 1 | 1 |
| HALLMARK_ESTROGEN_RESPONSE_LATE | 0.57399<br>32277 | 1 | 0.9926243568 |
| HALLMARK_INTERFERON_ALPHA_RESPONSE | -1 | 1 | 0.4714285714 |
| Interferon_alpha_beta_response | -1 | 1 | 0.4714285714 |
| Melanocytic | -<br>1.76629<br>5894 | 0 | 0.04285714286 |

**Table S13:** Liu RNA-seq - PFS Hazard correlation GSEA.

| Term | NES | NOM p-val | FDR q-val |
| --- | --- | --- | --- |
| Melanocytic | 2.213466259 | 0 | 0 |
| Mesenchymal_like | 1.904029631 | 0 | 0 |
| HALLMARK_EPITHELIAL_MESENCHYMAL_TRANSITION | 1.790718354 | 0 | 0.002272169 |
| HALLMARK_MYOGENESIS | 1.720209784 | 0 | 0.00170412675 |
| HALLMARK_HEDGEHOG_SIGNALING | 1.6081898 | 0.01351351351 | 0.013633014 |
| HALLMARK_NOTCH_SIGNALING | 1.524897754 | 0.04 | 0.040899042 |
| Mitochondrial(low_quality) | 1.476126272 | 0.01098901099 | 0.05258448258 |
| HALLMARK_ANGIOGENESIS | 1.457094219 | 0.06666666667 | 0.05708824613 |
| HALLMARK_TGF_BETA_SIGNALING | 1.449119138 | 0.03658536585 | 0.05983378367 |
| Neural_Crest_like | 1.445136148 | 0 | 0.05657700811 |
| HALLMARK_APICAL_JUNCTION | 1.441001886 | 0.0101010101 | 0.05267300864 |
| HALLMARK_UV_RESPONSE_DN | 1.413096972 | 0.02197802198 | 0.06134856301 |
| HALLMARK_HYPOXIA | 1.254733947 | 0.07142857143 | 0.2647950796 |
| HALLMARK_ESTROGEN_RESPONSE_EARLY | 1.244414099 | 0.1030927835 | 0.2687651332 |
| HALLMARK_ESTROGEN_RESPONSE_LATE | 1.236628976 | 0.08421052632 | 0.2717514124 |
| HALLMARK_UV_RESPONSE_UP | 1.206781854 | 0.1030927835 | 0.3208018607 |
| HALLMARK_GLYCOLYSIS | 1.177511306 | 0.1632653061 | 0.369294291 |
| HALLMARK_TNFA_SIGNALING_VIA_NFKB | 1.132847732 | 0.2 | 0.4782915746 |
| Stress (hypoxia response) | 1.129094043 | 0.2395833333 | 0.4638812396 |
| HALLMARK_KRAS_SIGNALING_DN | 1.121914217 | 0.2258064516 | 0.4607958732 |
| HALLMARK_UNFOLDED_PROTEIN_RESPONSE | 1.075012183 | 0.3448275862 | 0.5865441976 |
| HALLMARK_ANDROGEN_RESPONSE | 1.052356759 | 0.402173913 | 0.6305268976 |
| HALLMARK_KRAS_SIGNALING_UP | 0.993216495 | 0.54 | 0.8025696069 |
| HALLMARK_MYC_TARGETS_V1 | 0.9837268026 | 0.5416666667 | 0.7969632768 |
| HALLMARK_WNT_BETA_CATENIN_SIGNALING | 0.9512095122 | 0.5584415584 | 0.8651510685 |
| HALLMARK_CHOLESTEROL_HOMEOSTASIS | 0.9031004612 | 0.5227272727 | 0.9721387676 |
| HALLMARK_P53_PATHWAY | 0.8942327576 | 0.6804123711 | 0.9568356123 |
| HALLMARK_PANCREAS_BETA_CELLS | 0.8846672913 | 0.7341772152 | 0.9457903464 |
| HALLMARK_SPERMATOGENESIS | 0.8674471346 | 0.7765957447 | 0.9517254085 |
| Antigen_presentation | 0.8640009481 | 0.78125 | 0.927272169 |
| HALLMARK_APICAL_SURFACE | 0.8608831295 | 0.6891891892 | 0.9041766705 |
| HALLMARK_XENOBIOTIC_METABOLISM | 0.8545814536 | 0.79 | 0.8872109893 |
| HALLMARK_INFLAMMATORY_RESPONSE | 0.841169829 | 0.8 | 0.8813950113 |
| HALLMARK_FATTY_ACID_METABOLISM | 0.833323134 | 0.8041237113 | 0.8699065846 |
| HALLMARK_PEROXISOME | 0.8061126569 | 0.7912087912 | 0.8935466891 |
| HALLMARK_IL6_JAK_STAT3_SIGNALING | 0.7478988079 | 0.8488372093 | 0.9421927454 |
| HALLMARK_APOPTOSIS | 0.7239216134 | 0.9347826087 | 0.9392041268 |
| HALLMARK_MYC_TARGETS_V2 | -0.7030664514 | 0.9473684211 | 0.9766949153 |
| Mitotic | -1 | 1 | 0.4949556094 |
| HALLMARK_PROTEIN_SECRETION | -<br>1.00008618<br>1 | 0.5 | 0.51504237<br>29 |
| HALLMARK_BILE_ACID_METABOLISM | -<br>1.02730067<br>8 | 0.33333333<br>33 | 0.43911685<br>99 |
| HALLMARK_COAGULATION | -<br>1.03101353<br>8 | 0.25 | 0.44797551<br>79 |
| HALLMARK_IL2_STAT5_SIGNALING | -<br>1.05670393<br>2 | 0.33333333<br>33 | 0.40852442<br>67 |
| HALLMARK_DNA_REPAIR | -<br>1.07001397<br>6 | 0.2 | 0.39036016<br>95 |
| HALLMARK_MITOTIC_SPINDLE | -<br>1.07454145<br>9 | 0.5 | 0.40706214<br>69 |
| HALLMARK_OXIDATIVE_PHOSPHORYLATION | -<br>1.08715938 | 0.25 | 0.39618644<br>07 |
| Stress (p53 response) | -<br>1.10942787<br>5 | 0.26315789<br>47 | 0.36212516<br>3 |
| HALLMARK_ADIPOGENESIS | -<br>1.13638372<br>8 | 0.16666666<br>67 | 0.32627118<br>64 |
| HALLMARK_REACTIVE_OXYGEN_SPECIES_PATHWAY | -<br>1.26138463<br>2 | 0.08695652<br>174 | 0.12288135<br>59 |
| HALLMARK_PI3K_AKT_MTOR_SIGNALING | -<br>1.33451566<br>8 | 0 | 0.08389830<br>508 |
| HALLMARK_COMPLEMENT | -<br>1.37378046<br>6 | 0 | 0.05696798<br>493 |
| HALLMARK_G2M_CHECKPOINT | -<br>1.39059761<br>7 | 0 | 0.06408898<br>305 |
| HALLMARK_MTORC1_SIGNALING | -<br>1.42503813<br>5 | 0 | 0.06658595<br>642 |
| HALLMARK_HEME_METABOLISM | -<br>1.44594063<br>9 | 0 | 0.06991525<br>424 |
| HALLMARK_INTERFERON_ALPHA_RESPONSE | -<br>1.57141716<br>8 | 0 | 0.02796610<br>169 |
| Interferon_alpha_beta_response | -<br>1.59429249<br>5 | 0 | 0.01165254<br>237 |
| HALLMARK_E2F_TARGETS | -<br>1.65583943<br>2 | 0 | 0 |
| HALLMARK_INTERFERON_GAMMA_RESPONSE | -<br>1.71910937<br>2 | 0 | 0 |
| HALLMARK_ALLOGRAFT_REJECTION | -<br>1.98036875<br>7 | 0 | 0 |

**Table S1. (separate file)**

Primary screen asgRNA abundances Create a page break and paste in the table above the caption.

**Table S2. (separate file)**

Primary screen ICB vs NSG log fold changes and hit calls

**Table S3. (separate file)**

Secondary screen asgRNA abundances

**Table S4. (separate file)**

Secondary screen ICB vs NSG log fold changes and hit calls

**Table S5. (separate file)**

Bulk RNA-seq differential expression CREB5 sg2 vs CTRL

**Table S7. (separate file)**

Bulk RNA-seq differential expression CREB5 sg1 vs CTRL

**Table S8. (separate file)**

Bulk RNA-seq CREB5 sg1 vs CTRL GSEA results

**Table S11. (separate file)**

Jerby-Anon scRNA-seq - TF correlation with Mesenchymal

**Table S14. (separate file)**

Patients’ demographics

